# The rat frontal cortex encodes a value map for economic decisions under risk

**DOI:** 10.1101/2021.11.19.469107

**Authors:** Xiaoyue Zhu, Chaofei Bao, Joshua Moller-Mara, Jingjie Li, Sylvain Dubroqua, Jeffrey C. Erlich

**Author notes:** Indicates equal contribution to the manuscript (Xiaoyue Zhu, Chaofei Bao).

## Abstract

Neurons in frontal and parietal cortex encode task variables during decision-making, but causal manipulations of the two regions produce strikingly different results. For example, silencing the posterior parietal cortex (PPC) in rats and monkeys produces minimal effects in perceptual decisions requiring integration of sensory evidence, but silencing frontal cortex profoundly impairs the same decisions. Here, we tested the causal roles of the rat frontal orienting field (FOF) and PPC in economic choice under risk. On each trial, rats chose between a lottery and a small but guaranteed surebet. The magnitude of the lottery was independently varied across trials and was indicated to the rat by the pitch of an auditory cue. As in perceptual decisions, both unilateral and bilateral PPC muscimol inactivations produced minimal effects. FOF inactivations produced substantial changes in behavior even though our task had no working memory component. We quantified control and bilateral inactivation behavior with a multi-agent model consisting of a mixture of a ‘rational’ utility-maximizing agent (*U* = *V^ρ^*) with two ‘habitual’ agents that either choose surebet or lottery. Effects of FOF silencing were best accounted for by a decrease in *ρ*, the exponent of the utility function. The decrease in *ρ* could be parsimoniously explained by a dynamical model where the FOF is part of a network that represents the value of the lottery to compare against a remembered boundary representing the value of the surebet. This dynamical model predicted that neurons in the FOF should increase their activity with increasing lottery values. In line with our predictions, single neurons recorded in the FOF encoded the value of the lottery even when controlling for choice. These results together demonstrate that FOF is a critical node in the neural circuit for decision under risk.

## Introduction

Understanding decisions under risk is of substantial interest from a public health and welfare perspective: excessive risk-taking is associated with drug and gambling addiction^1^, dangerous teen driving^2^ and other pathologies^3^. On the other hand, inadequate risk-taking is also undesirable: people who avoid investing in the stock market can have their savings diminished by inflation; a mouse that is unwilling to risk predation for foraging will starve. Data from twin and genome-wide association studies^4–7^ suggest that genetics accounts for ~ 30% of variation in risk-tolerance, indicating that animal models can help establish the link between genes, brains and risk-tolerance.

The neurobiology of risky decision-making has been studied in human, non-human primate and rodent subjects reviewed in^8–10^. Rodent work has mostly focused on decision-making under uncertainty in new or changing environments, that is, the neural mechanisms for learning the values of actions. Action-value learning under ‘unexpected uncertainty’ has also been studied extensively in monkeys and humans^11–13^. These studies typically identify regions that are associated with learning action-values in general: amygdala^14–16^, basal ganglia^17–19^, and orbital and medial prefrontal cortex^20–22^. Human, monkey, and, to a lesser extent, rodent work has also examined the neurobiology of decision-making under risk when the probabilities are known, i.e., ‘expected uncertainty’^23–27^. This is closer to the way risky decisions are studied in economics or finance research, where the potential outcomes of different actions are given explicitly on each trial. In these studies, activity in regions associated with orienting decisions^28^ including the parietal cortex^10,29,30^ and frontal cortex^31,32^ represents the value of the options, as the subjects were typically asked to respond by shifting gaze to a spatial target.

One challenge in synthesizing the vast literature on decisions under risk is that risk-tolerance is not monolithic^33^. When behavior is measured either in the laboratory or in real-life, any avoidance of uncertainty can be considered as ‘risk-aversion’, but such avoidance can come from distinct cognitive constructs. For example, risky behavior in teenagers may result from an incomplete perception of risk associated with those behaviors rather than a greater tolerance for the actual risk ^34^. Reinforcement learning agents with high learning rates will seem more risk-averse because they will avoid actions after a single loss (i.e. ‘lose-shift’), even if on average that action provides good outcomes ^35,36^. Under the expected utility framework, risk-aversion is usually associated with a decelerating utility function: the more rapid the deceleration, the more risk-averse the subject^37–39^. In finance, risk-aversion is typically modeled as variance-aversion^40^. This rich taxonomy of constructs underlying risk-preference not only adds confusion when parsing the literature, but also makes it challenging to the design of animal experiments which estimate all elements simultaneously.

Here, we present results from a risky choice task where the animals make choices under ‘expected uncertainty’ on a trial-by-trial basis. On each trial, rats made decisions between a ‘surebet’ (small but guaranteed reward) and a lottery with fixed probability and cue-guided magnitude. Our model-based quantification of animals’ behavior incorporated parameters to capture marginal utility, decision noise, and choice biases. This task and modeling framework provided a foundation for rigorous decomposition of neurobiological elements of risky choice. With this framework, we examined the causal contribution of the frontal orienting field (FOF) in frontal cortex and the posterior parietal cortex (PPC): two cortical areas that have been implicated in perceptual decision-making^28,41–43^.

In perceptual decision that require working-memory, the FOF seems to be essential for maintaining a plan of the upcoming choice. Unilateral silencing of FOF biases animals towards the ipsilateral choice and this bias is larger for trials with longer memory periods^44^. Bilateral silencing also generates an impairment that grows with longer delays or periods of integration ^41^. In order to distinguish the role of working-memory from the cognitive processes required for economic choice under risk (i.e. trading off the cost of uncertainty with the benefits of a larger reward), we did not include a working-memory component in our task. Nonetheless, we predicted that unilateral silencing of FOF would cause contralateral impairments in economic choices, due to our hypothesis that FOF serves as a bottleneck for higher order cognitive processes to guide orienting decisions. For bilateral FOF the prediction was less clear: we expected it to influence behavior, possibly by increasing the decision noise.

In contrast to the results from FOF, we previously found that unilateral silencing of PPC in rats did not bias perceptual decisions. PPC only biased ‘free-choice’ trials, where animals were rewarded regardless of the left or right response^41^. We speculated that the difference between the efficacy of PPC inactivations in perceptual vs. free choice might be that PPC only plays a causal role when decisions are internally guided. Risky choices are internally guided in the sense that each subject has some risk-preference: there is no single ‘correct answer’ on each trial. Moreover, signatures of expected value are reliably found in PPC^29,30^. Thus, we hypothesized that PPC silencing might influence economic choices, in contract to perceptual decisions.

Although there is substantial literature comparing different functional forms of decision under risk^45–47^, we are unaware of any previous studies that simultaneously estimates these parameters, and examines how silencing of frontal or parietal cortices shifts specific cognitive constructs underlying risky choice. We found that PPC silencing had minimal effects on decisions under risk (but biased free choice). Surprisingly, we found that bilateral silencing of FOF shifted animals away from choosing the lottery. Model-based analysis of these results indicated that the shift was likely caused by a change in the utility function. Moreover, this effect was parsimoniously explained by a dynamical model where the FOF as a part of a network for encoding the value of the lottery. This dynamical model predicted that the FOF should contain neurons that monotonically increase with the expected value of the lottery. To test this, we recorded neurons in the FOF and found that a substantial portion of them encoded the expected value of the lottery, even when controlling for the animals’ choice. Together, these results suggests that the FOF is a key node in a network for encoding a value map in the service of economic choice.

## Materials and Methods

### Subjects

A total of 14 rats (13 males, 1 female) were used in this study, including 12 Sprague Dawley rats and 2 male Brown Norway rats (Vital River, Beijing, China). Animals were paired housed during the training period and then single housed after electrode or cannula implantation. Of these 14 animals, six male Sprague Dawley rats and two male Brown Norway rats were used for FOF/PPC muscimol inhibition experiment. These 8 animals were placed on a controlled-water schedule and had access to free water 20 minutes each day in addition to the water they earned in the task. The other six Sprague Dawley rats (1 female) were used for the *in vivo* electrophysiology recording experiment. For these animals, 4% citric acid water was available *ad libitum* in their home cage instead of controlled-water-access. All rats were kept on a reversed 12 hour light-dark cycle and were trained during their dark cycle. Animal use procedures were approved by New York University Shanghai International Animal Care and Use Committee following both US and Chinese regulations.

### Behavioral Apparatus

Animal training took place in custom behavioral chambers, located inside sound- and light-attenuating boxes. Each chamber (23 × 23 × 23 cm) was fit with 8 nose ports arranged in four rows (Figure 1A), with speakers located on the left and right side. Each nose port contained a pair of blue and a pair of yellow light emitting diodes (LED) for delivering visual stimuli, as well as an infrared LED and infrared phototransistor for detecting rats’ interactions with the port. The port in the bottom row contained a stainless steel tube for delivering water rewards. In the risky choice task, only 4 of the 8 ports were used. Other tasks in the lab utilized all 8 ports. Each training session lasted for around 90 minutes.

**Figure 1.**
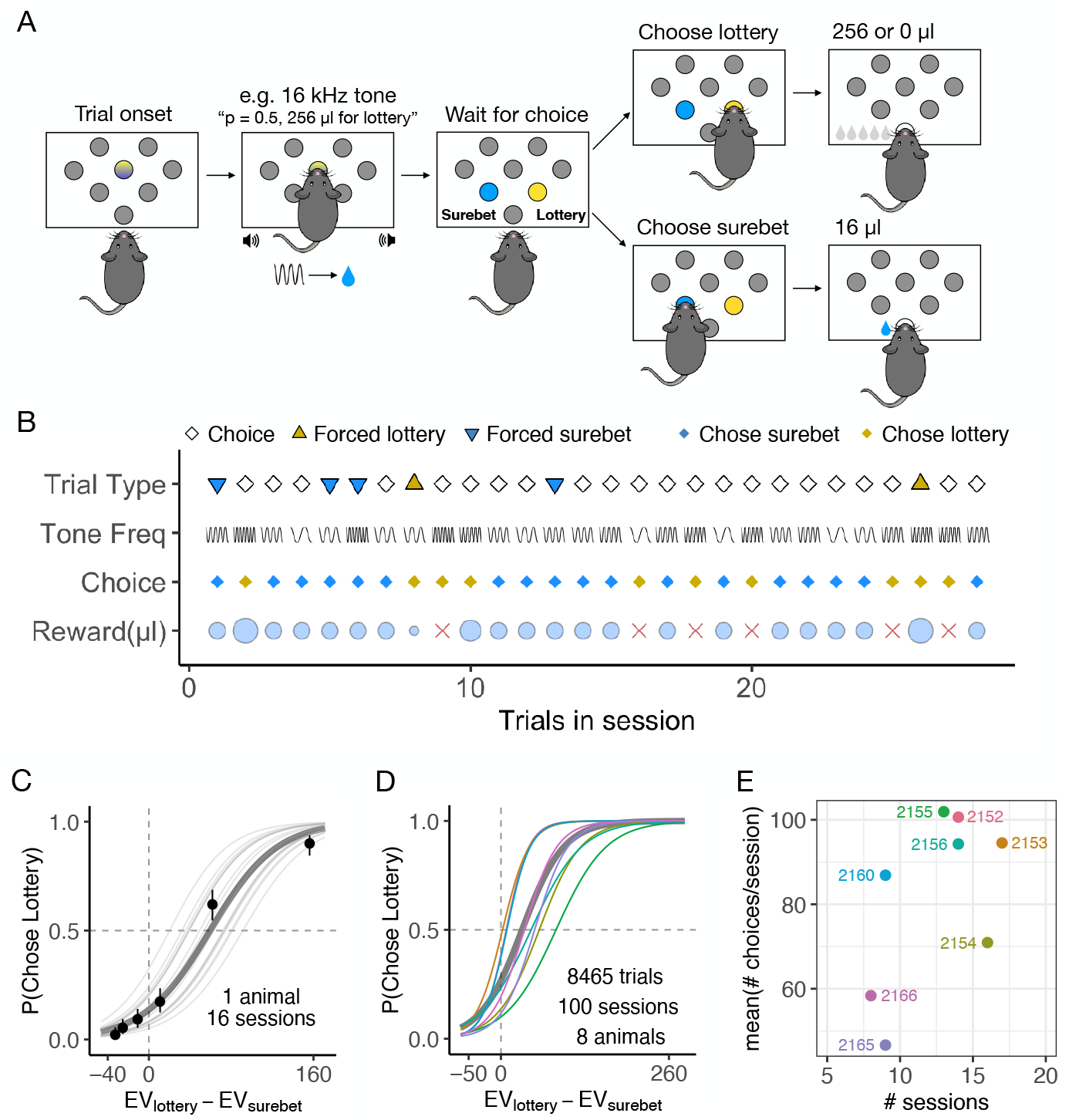
The risky choice task and animal behavior. **A.** Schematic of the risky choice task. Each trial began with the onset of central LED, which cued the animal to poke into the center port and hold there for 1 s. A tone was played, indicating the magnitude of the lottery. After 1 s, the animal withdrew from the center port and made a choice poke into the surebet or the lottery port. The lottery sound was played until the choice poke. A surebet choice delivered a small and guaranteed reward, whereas a lottery poke gave either 0 or the cued magnitude based on lottery probability, which was fixed for each animal. See more detailed task description in Methods. **B.** Timeline of trials in one example session. For trial type, white diamond, yellow triangle and blue triangle represents the choice trial, forced lottery trial and forced surebet trial, respectively. The sine-waves in the ‘tone freq’ row indicate the lottery magnitudes cued by the tone frequency. The more compact the sine-wave is, higher is the lottery magnitude on this trial. Animal’s choices are marked in diamonds, with yellow for lottery and blue for surebet. The reward received (μL) on each trial is shown in light blue circles, whose size represents the relative amount. The red cross indicates a lost lottery with no rewards. **C.** Example subject performance from 16 control sessions 1 day before an infusion. The probability of choosing lottery is plotted as as a function of the expected value of lottery minus the expected value of surebet (*V_lottery_P_lottery_* – *V_surebet_*), where *V* represents *μ*L of water. The circles with error bars are the mean and 95% binomial confidence intervals. The lines are the psychometric curves generated by a logistic fit, the thin gray lines are fit to each session, the thick gray line fit to all the sessions combined. **D.** Subject performance from 100 control sessions 1 day before an infusion event (8 rats). The colored lines are the psychometric curves generated by a logistic fit to all the control sessions from each animal, the thick gray line fit to all the sessions combined. **E.** The number of control sessions within the infusion period, and the average number of choice trials, colored by subject. The text indicates the subject ID.

### Behavior

Trials began with both yellow and blue LED turning on in the center port. This cued the animal to poke its nose into the center port and hold it there for 1 s, after which the center lights were turned off and the choice ports became illuminated. We refer to this period as the ‘soft fixation’ period, as the animal was allowed to withdraw any time after the initial poke. From here, if the animal poked into a different port other than the center port, a short white noise would play to indicate that this is a mistake. The choice ports were triggered as soon as the animal performed a second poke into the center port. All animals exploited the soft fixation strategy, albeit to different degrees individually. They tended to withdraw after the initial poke but stayed close to the center port during the soft fixation period (Figure S1B).

During the soft fixation period a tone played from both speakers, indicating the lottery magnitude for that trial. There were 6 distinct frequencies indicating different lottery magnitudes (2.5 kHz – 20 kHz, 75 dB). The frequency of each lottery was around one octave away from the adjacent tones, making distinguishing the different offers perceptually easy^48^. The cue frequency-to-lottery magnitude mapping and the location of the surebet port were counterbalanced across animals. At the end of fixation, the lottery port and surebet port were illuminated with yellow and blue lights, respectively. The tone stopped as soon as the animal made a choice by poking into one of the choice ports. If the animal chose surebet, a small and guaranteed reward would be delivered at the reward port. If the animal chose lottery, it would either receive the corresponding lottery magnitude or nothing based on the lottery probability, which was titrated for each animal and ranged from 0.5 to 0.75 across all subjects. We refer to these trials as ‘choice’ trials. In order to ensure that the subjects experienced all the outcomes, the choice trials were randomly interleaved with trials that we refer to as ‘forced’ trials. The forced trials differ from choice trials in that only one of the two ports was illuminated and available for poking, forcing the animal to make that response. The forced surebet and forced lottery trials together accounted for 25% of the total trials. The inter-trial interval (ITI) was between 3 and 10 seconds (uniformly distributed). A trial was considered a violation if the animal failed to poke into the center port within 300 seconds from trial start, or it did not make a choice 30 seconds after fixation. Violations were excluded from all analyses, except where they are specifically mentioned.

In some sessions, ‘free’ trials were interleaved with the choice and forced trials. Free trials were similar to choice trials, and at the end of fixation both left and right port were illuminated with blue LEDs. The animal would receive a medium-sized reward (2 times the surebet) regardless of which port it chose. The free trials were randomly interleaved with the choice and forced trials, and were introduced only after all the experiments presented in Figure 2 & 3A-D were completed.

**Figure 2.**
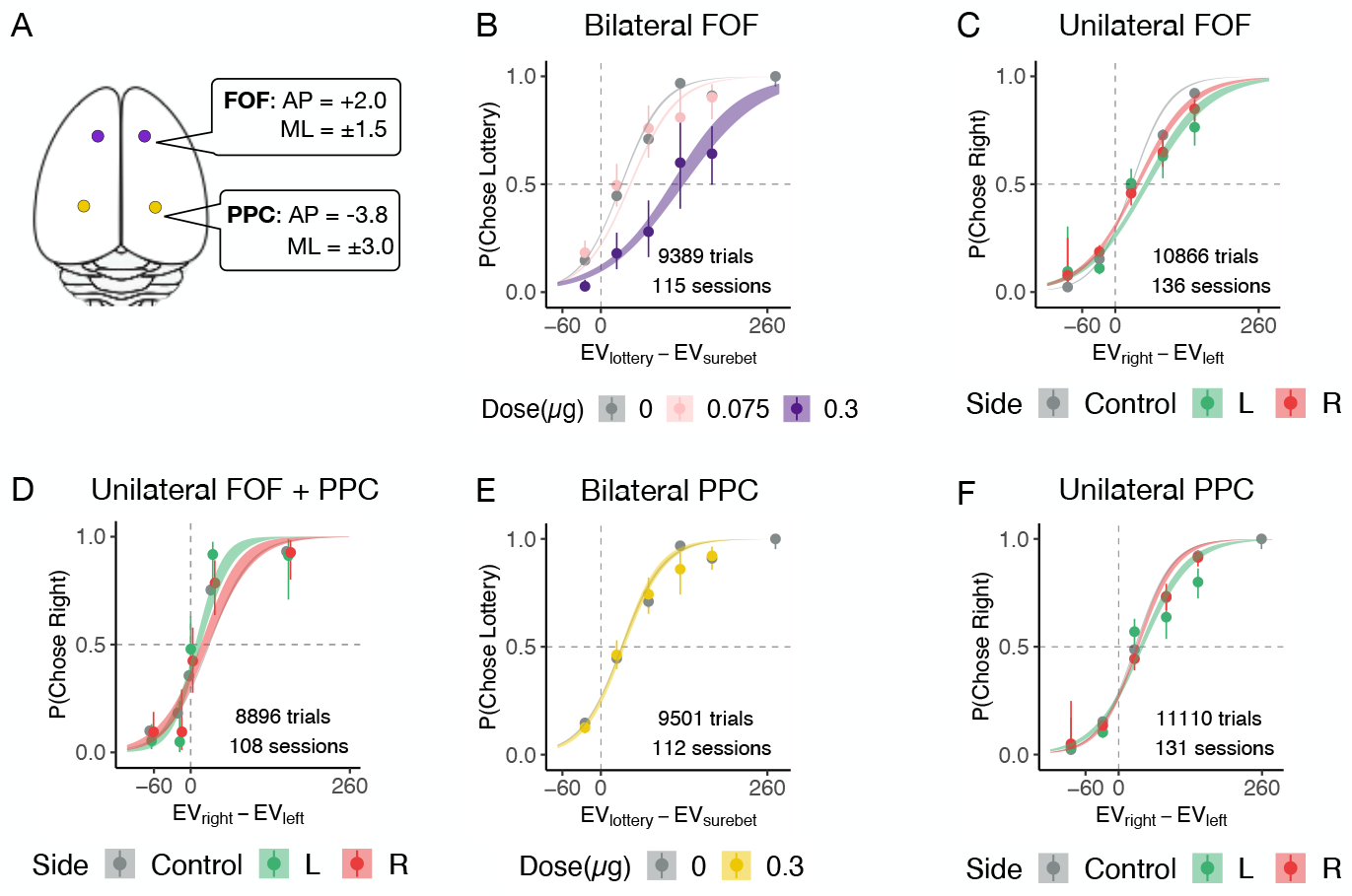
Bilateral and unilateral infusions in FOF and PPC. **A.** Top-down view of the rat cortex with the target coordinates of FOF and PPC, where the cannulae were implanted. **B.** Bilateral infusion of muscimol (0.3 *μ*g) into the FOF significantly shifted the choices towards the lottery. Control sessions 1 day before an infusion are shown in gray (n = 17 sessions, 8 rats), 0.075 *μ*g per side bilateral FOF infusions (n = 6 sessions, 5 rats) are in light pink, 0.3 *μ*g per side bilateral FOF infusions (n = 9 sessions, 8 rats) are in dark purple. The circles with error bars are the mean and 95% binomial confidence intervals. The ribbons are from a logistic fit to the data. See details in Methods. **C.** Unilateral infusion of muscimol into the left and right FOF resulted in a small but reliable shift towards surebet. Control sessions are in gray (n = 39 sessions, 8 rats), 0.3 *μ*g left FOF infusions (n = 16 sessions, 8 rats) are in green, 0.3 *μ*g right FOF infusions (n = 20 sessions, 8 rats) are in red. **D.** Simultaneous unilateral inactivation of FOF and PPC had no significant effect. Control sessions are in gray (n = 8 sessions, 4 rats), 0.3 *μ*g left FOF infusion and 0.6 *μ*g left PPC infusions (n = 4 sessions) are in green, 0.3 *μ*g right FOF infusion and 0.6 *μ*g right PPC infusion (n = 4 sessions) are in red. **E.** Bilateral infusion of muscimol into the PPC had no significant effect. Control sessions are in gray (n = 24 sessions, 7 rats), 0.3 *μ*g per side bilateral PPC infusions (n = 12 sessions, 7 rats) are in gold. **F.** Unilateral infusion of muscimol into the PPC had no significant effect. Control sessions are in gray (n = 31 sessions, 8 rats), 0.3 *μ*g left PPC infusions (n = 11 sessions, 7 rats, 3) are in green, 0.3 *μ*g right PPC infusions (n = 19 sessions, 8 rats) are in red.

**Figure 3.**
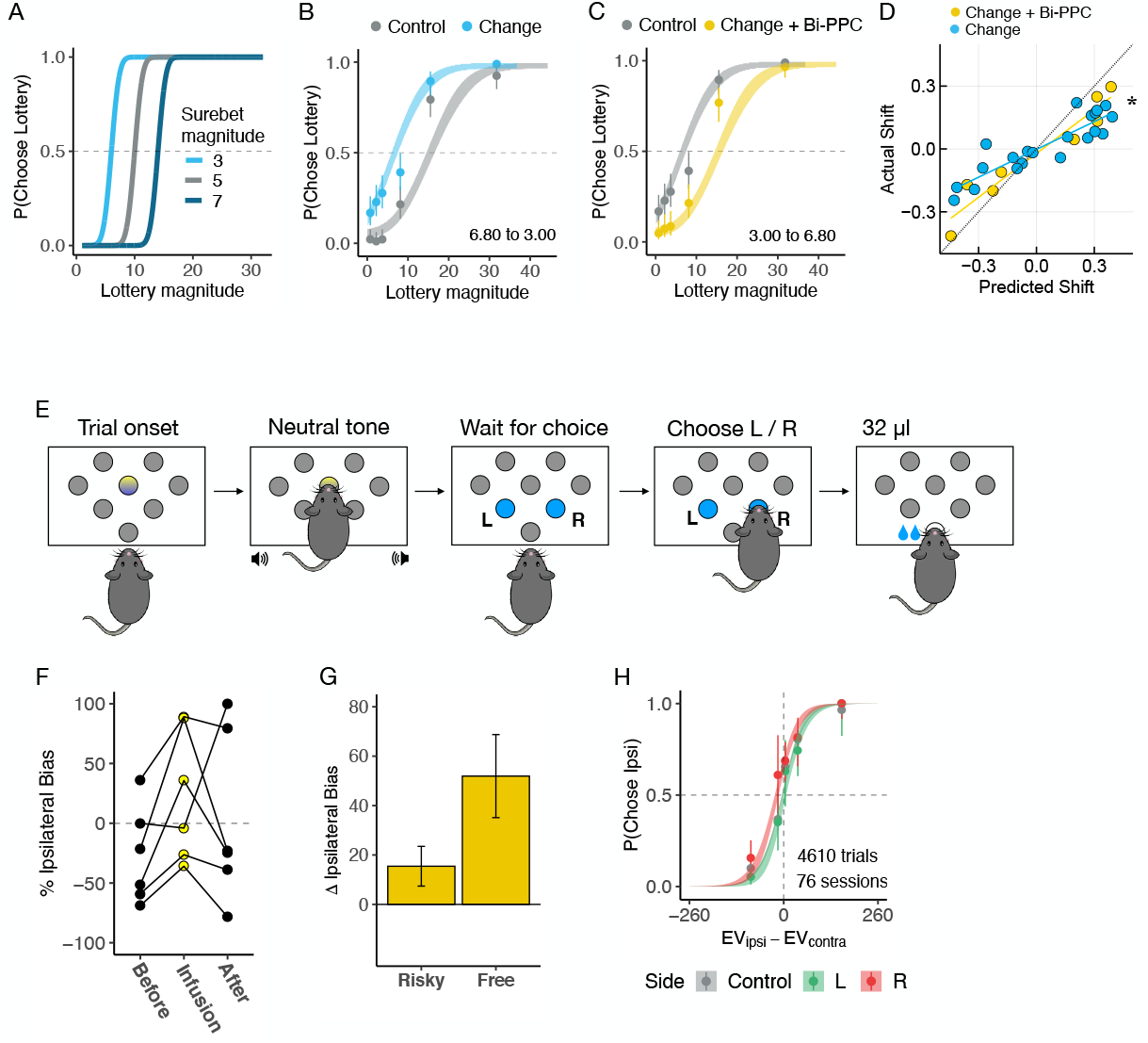
Further investigation into the potential roles of PPC. **A.** Schematic showing changing the surebet magnitude is equivalent to shifting the choice boundary. The data points were simulated from a risk-neutral agent using the three-agent model (*ρ* = 1, *σ* = 3, *ω_rational_* = 1). A smaller surebet magnitude (light blue) horizontally shifts the psychometric curve leftwards, a larger surebet magnitude (dark blue) shifts the curve rightwards. The frequency-to-lottery mapping remains the same. **B.** Changing surebet magnitude from 6.8 to 3 shifted choices leftwards in one example animal. Combined trials from 6 sessions before the change are shown in gray, after the change shown in blue. One three-agent model was fit to all the trials and the parameters were used for ribbon extrapolation. **C.** Same as B but with 0.6 *μ*g per side bilateral PPC infusion, performed on the day of surebet change (from 3 to 6.8). **D.** The three-agent mixture model predicts the shifts in behavior well. One model was fit using all the sessions containing various surebet magnitudes for each animal. On x-axis is the predicted shift in probability choosing lottery (*p*(Choose Lottery)): the difference in *p*(Choose Lottery) between model prediction using the new surebet magnitude and the session just before that change. On y-axis is the actual shift in *p*(Choose Lottery): the difference in *p*(Choose Lottery) between the first session of a surebet change and the session before that change. Sessions with just surebet change are in blue (n = 21; 4 animals), sessions with both surebet change and 0.6 *μ*g per side bilateral PPC infusions are in gold (n = 8). There is a strong correlation between predicted and actual shift, and this slope of this relationship is significantly different between shifts where PPC was silenced and control shifts. **E.** Schematic of the free trials. After fixation at the center port accompanied by a neutral tone, the animal was free to choose the left or right port, both illuminated in blue LEDs. Choosing either port resulted in a reward twice the magnitude of surebet. The free trials were randomly interleaved with the forced and choice trials. **F.** Unilateral PPC infusions (0.6 *μ*g) led to a significant ipsilateral bias towards the side of infusion. This panel shows % ipsilateral bias: ∑ choose_infusion_side – ∑ choose_other_side)total_choices, when the side of infusions was chosen to be the opposite to the animals’ preferred side. % ipsilateral bias was computed using free trials from the previous 3 sessions, the infusion session, and the following 3 sessions for 6 subjects. **G.** Unilateral PPC infusions generated a significant 52 ± 16% (mean ± s.e. across rats, n = 6) change in % ipsilateral bias on free trials compared to control sessions (3 pre-infusion sessions). For the choice trials from the same sessions, the change in % ipsilateral bias was not significant (15 ± 8%). **H.** Performance on the choice trials was not affected. Control sessions from the 3 pre-infusion sessions (n = 65 sessions, 6 rats) are in gray, 0.6 *μ*g left PPC infusions (n = 5 sessions) are in green, 0.6 *μ*g right PPC infusions (n = 6 sessions) are in red.

An additional stage was added to the electrophysiological recording animals to gradually shift the ‘soft fixation’ to a ‘hard fixation’. In this stage, trials began with a 300ms ‘hard fixation’ and followed with a 700ms ‘soft fixation’. The rats must hold their nose into the central port during the ‘hard fixation’ time. If the rats failed to maintain the central nosepoke during the ‘hard fixation’ time, the fixation timer stopped and the trial restart to let the animal attempt again immediately. The ‘hard fixation’ period adaptively increased while the ‘soft fixation’ period decreased, until the entire fixation period became ‘hard fixation’.

#### Training pipeline

Animal training took place in two distinct phases: the operant conditioning phase and the risky choice phase. Briefly, in the operant conditioning phase, rats became familiar with the training apparatus and learned to poke into the reward port when illuminated. Trials began with the illumination of the reward port, and water reward was immediately delivered upon port entry. After the rats learned to poke in the reward port reliably, they proceeded to the next training stage where they had to first poke into an illuminated choice port (left or right, chosen randomly) before the reward port was illuminated for reward. They graduated to the risky choice phase if they correctly performed these trials at least 40% of the session.

In the risky choice phase, rats started with only two frequencies: the lowest and highest, corresponding to the smallest and largest lottery magnitude. Initially, there were more forced trials than choice trials to help them understand the task. Once the animals reliably differentiated between the low and high lottery choice trials, we increases the ratio of choice trials to force trials. Intermediate frequencies were added one by one, contingent upon good behavior in the choice trials with existing frequencies. The lottery probability and the surebet magnitude were adapted to each animal so that their preferences could be reliably estimated. For example, if an animal chose the lottery too often, the lottery probability would be decreased. The goal was to be able to accurately estimate parameters of the three-agent mixture-model (described below).

### Surgery for intracranial muscimol infusion

Surgical methods were similar to those described in Erlich et al. ^41^. The rats were anesthetized with isoflurane and placed in a stereotaxic apparatus (RWD Life Science Co.,LTD, Shenzhen). The scalp was shaved, washed with ethanol and iodopovidone and incised. Then, the skull was cleaned of tissue and blood. The stereotax was used to mark the locations of craniotomies for the left and right FOF and PPC, relative to Bregma on the skull. Four craniotomies and durotomies were performed and the skull was coated with a thin layer of C&B Metabond (Parkell Inc., NY). Each guide cannula along with the injector (RWD Life Science Co.,LTD, Shenzhen) was inserted 1.5 mm into the cortex measured from brain surface for each craniotomy. The guide cannulae were placed and secured to the skull one at a time with a small amount of Absolute Dentin (Parkell Inc., NY). The injector was removed from each guide once the guide was secured to the skull. After all four guide cannulae were in place, more Absolute Dentin was applied to cover the skull and further secure the guide cannulae. Vetbond (3M, U.S.) was applied to glue the surrounding tissue to Absolute Dentin. The animals were given 7 days to recover on free water before resuming training.

### Cannulae

All 8 rats were implanted bilaterally in FOF (+2 AP, ±1.5 ML mm from Bregma) with 26 AWG guide cannulae (RWD Life Science Co.,LTD, Shenzhen) and in lateral PPC (−3.8 AP, ±3.0 ML from Bregma) with 26 AWG guide cannulae (4 cannulae per rat total). The tip of the guide sat on the brain surface, while the 33 AWG injector was extended 1.5 mm below the bottom of the guide cannula. Dummy cannula (which were left in the guides in between infusions) extended 0.5 mm past the guides into the cortex. Cannula placement were verified post-mortem (Figure S2B-E).

### Infusions

Infusions were performed once a week with normal training days taking place on all other days. This was to minimize adaptation to the effects of the muscimol and to have stable performance in the sessions immediately before infusion sessions. Animals were held by an experimenter during the infusion, no general anesthetic was administered. On an infusion day, the rat was placed on the experimenter’s lap and the dummy cannulae were gently removed and cleaned with iodine and alcohol and then rinsed in DI water. The injector was inserted into the target guide cannula and reached 1.5 mm into cortex. A 1 *μL* syringe (Gaoge, Shanghai) connected via tubing filled with mineral oil to the injector was used to infuse 0.3 *μ*L of muscimol (of various concentrations) into cortex. The injection was done over 1 minute, after which the injector was left in the brain for 5 more minutes to allow diffusion before removal. The thoroughly cleaned and rinsed dummies were placed into the guide cannula. The rats began training 2 to 53 minutes after the infusion, the average time between infusion and starting of the behavioral session was 27 minutes. See Figure S2 for the complete list of all infusion doses, regions, and order for each rat.

### Data analysis for the behavior and muscimol infusions

For all analyses, we excluded time out violation trials (where the subjects disengaged from the ports for more than 30 s during the trial) and trials with reaction time longer than 3 s. Unless otherwise specified, the ‘control’ sessions refer to the sessions one day before any infusion event during the course of the experiment. All analysis and statistics for the infusion experiments were computed in R (version 3.6.3, R Foundation for Statistical Computing, Vienna, Austria).

#### Generalized linear mixed-effects models (GLMM)

Generalized-linear mixed-effects models (GLMM) were fit using the lme4 R package^49^. To test whether bilateral infusion had any effects on performance, we specified a mixed-effects model where the probability of a lottery choice was a logistic function of *EV_lottery_* – *EV_surebet_*, muscimol dosage (μg) and their interaction as fixed effects. The rat and an interaction of rat, *EV_lottery_* – *EV_surebet_* and dosage were modelled as within-subject random effects. For this model (and the other GLMM described in this session) the control sessions were coded as a 0 *μ*g dose. The expected value of lottery is the product of the lottery probability and lottery magnitude (*EV_lottery_* = *P_lottery_ · V_lottery_*). Similarly, *EV_surebet_* denotes the expected value of surebet, which is simply the value of surebet here (*EV_surebet_* = *V_surebet_*, since *P_surebet_* = 1). In GLMM formula syntax:

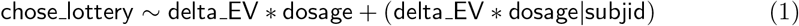

where chose_lottery is 1 if lottery was chosen on a trial, delta_EV is *EV_lottery_ – EV_surebet_* and subjid is the subject ID for each rat.

To test whether unilateral infusions caused a left/right bias as in^41^, we specified a mixed-effects model similar to the one described above:

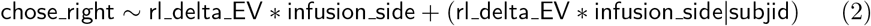

where chose_right is 1 if the right port is chosen on this trial, rl_delta_EV is *EV_right_* – *EV_left_* and infusion_side is a categorical variable with three levels: left, right and control. The plots in Figure S3 show that the model fits for each rat, reflecting how the random effects allow for each rats’ data to be fit, while also evaluating the significance of fixed effects.

To estimate the shift in indifference point induced by bilateral FOF inactivation, we first fit a GLMM as described above. We generated synthetic data points for delta_EV to extend its range, and the model was used to predict *p*(Choose Lottery) for each synthetic data point. For each animal, we identified the delta_EV values that resulted in *p*(Choose Lottery) to be between 0.499 and 0.501, which is the definition of indifference point. The average indifference point was obtained by taking the mean of such values across animals.

To test whether unilateral PPC infusions led to an ipsilateral bias in both free choice and risk choice trials, we specified a GLMM as following:

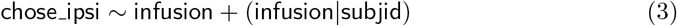

where chose_ipsi is a binary variable indicating whether the animal chose the side ipsilateral to the infusion side or not, and infusion is a binary variable representing the presence of an unilateral PPC infusion.

To estimate changes in reaction time, we used Linear Mixed-Effects Models (LMM). The formula for bilateral infusion was:

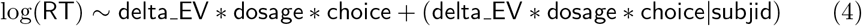

where log(RT) denotes the logarithm of reaction time, choice is a binary value for the surebet/lottery choice (0/1). Similarly, the formula for unilateral infusion was:

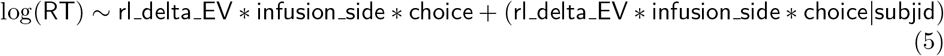

To test whether the outcome of the previous trial affected choice on the current trial, we first classified the previous trial’s outcome into three categories: lottery-win, lottery-lose and surebet. If the previous trial was a violation, we considered that as a surebet choice.

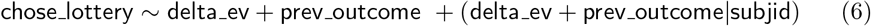

where prev_outcome is a categorical variable with three levels of previous outcome as above.

#### Surebet learning

To test the role of PPC in learning, we periodically changed the surebet magnitude in a model-based way to shift the decision boundary. For each shift, we fit the three-agent model (described below) on control data from the past 14 days to obtain a set of parameters. Using a binary search algorithm, we then used those parameters to generate synthetic choices with different surebet magnitudes until we found a value that produced a shift in probability choosing lottery (*p*(Choose Lottery)) close to the target (drawn uniformly from ±*U*(0.2, 0.3)). The new surebet magnitude was assigned to the animal on the day of change. All animals in the surebet learning experiment had undergone two rounds of shift without any infusion, in the course of 14 days, to acclimate them to the new routine before bilateral PPC infusions. The first two surebet change sessions are not included in the analysis of Figure 3.

To test whether bilateral PPC infusions (0.6 mg/kg) changed the slope of the actual shift in response to a predicted shift we fit a linear model

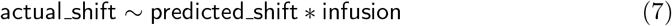

and reported the statistics of the interaction term, *β_predicted_shift:infusion_*

#### The three-agent mixture model

We developed a three-agent mixture model that used 4 parameters to transform the offers on each trial into a probability of choosing lottery as a weighted outcome of three agents (Figure 4A): a rational agent, a ‘lottery agent’ and a ‘surebet’ agent. For the rational agent, we assume an exponential term *ρ* for the utility function, *U* = *V^ρ^*. A concave utility function (*ρ* < 1) implies risk aversion, a linear function with *ρ* =1 implies being risk-neutral and a convex function (*ρ* > 1) implies risk seeking. We captured stochasticity in the animals behavior modeling the internal representation of expected utility as a Gaussian random variable.

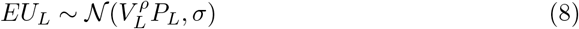

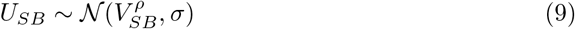

where the expected utility of lottery, *EU_L_*, and the utility of the surebet, *U_SB_* are Normal distributions. *V_L_, V_SB_* refer to the magnitude of lottery and surebet and *P_L_* is the probability of lottery payout. The probability of choosing lottery for the rational agent then becomes

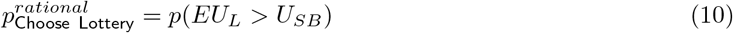

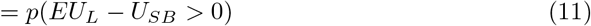

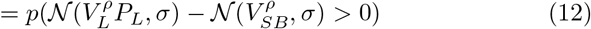

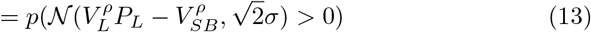

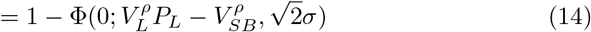

where 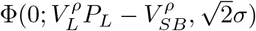 is the cumulative Normal distribution with mean 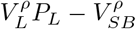, standard deviation 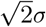 and evaluated at 0. Note that this provides fits with similar likelihood as the softmax choice function with *β* as temperature:

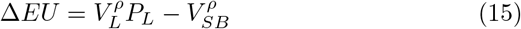

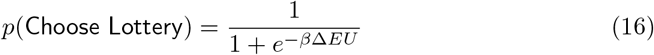

The other two agents in the three-agent mixture model are the lottery and surebet agents. They represent the habitual bias of the animal to make one or the other choice regardless of the lottery offer, similar to biased lapse terms in Erlich et al. ^41^. The probability of choosing lottery for the lottery agent is 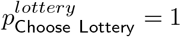 and for the 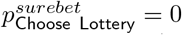.

**Figure 4.**
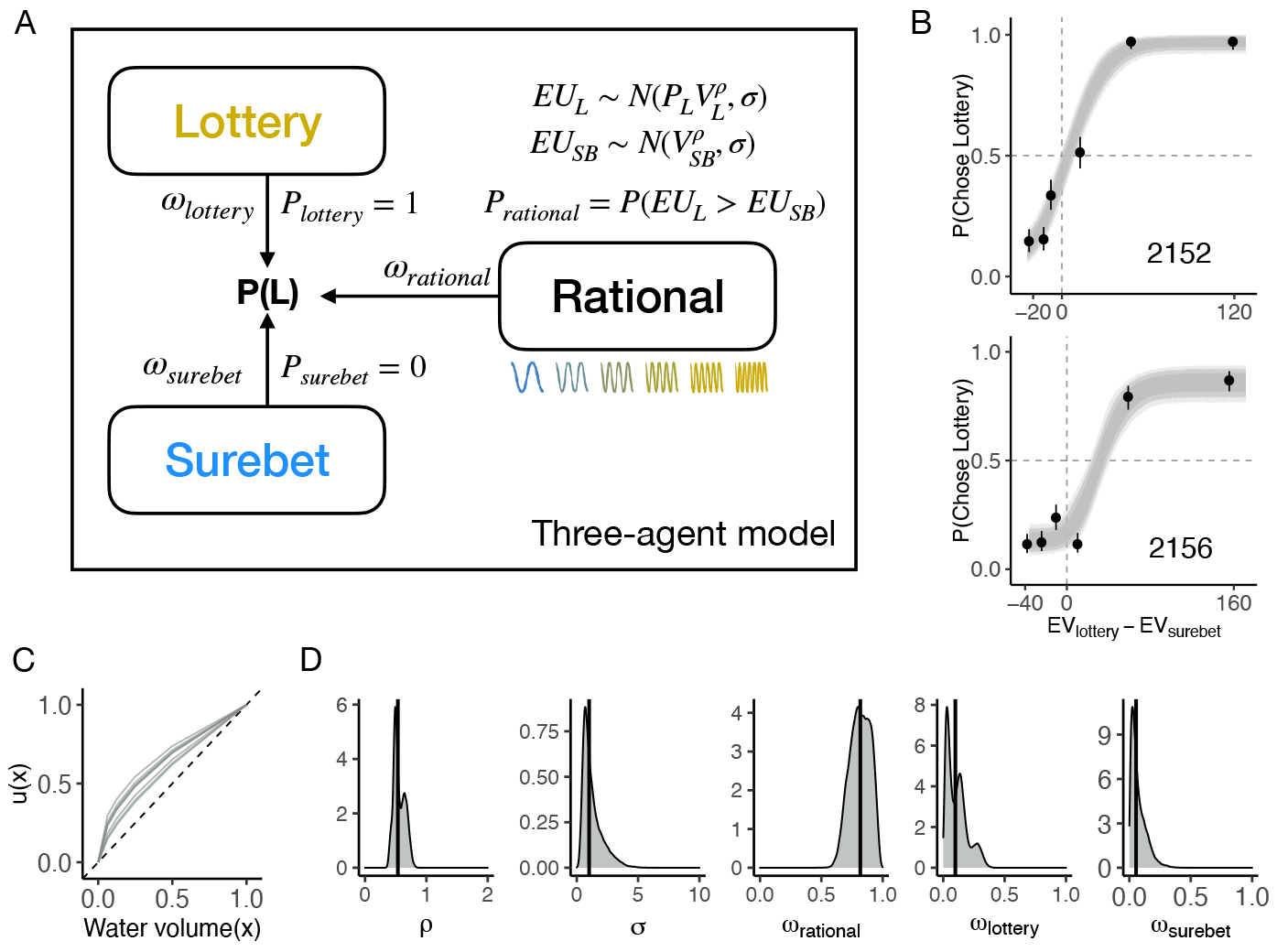
The three-agent mixture model and model fits. **A.** The three-agent mixture model. The animal’s choice is modelled as a weighted average of the three agents, each implementing a different behavioral strategy to perform the task. Each agent outputs a probability of choosing lottery that makes up the probability vector 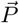, which is combined using their respective weights 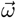. See Methods for model details. **B.** The three-agent mixture model can fit the control behavior well. The circles with error bars are the binned mean and 95% binomial confidence intervals. The ribbons are model predictions generated using the fitted parameters. The dark, medium and light shade represent 80%, 95% and 99% confidence intervals, respectively. **C.** The subjective utility functions for each rat computed using maximum a posteriori *ρ* estimation, normalized by the maximum water volume. **D.** Density plots of concatenated posterior samples (4000 each) from the model fits across 8 animals. The black bar is the median of the distribution.

The last step is to obtain *p*(Choose Lottery) by mixing the probability from each agent 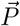 with their respective mixing weights 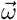 that sum up to 1. Formally,

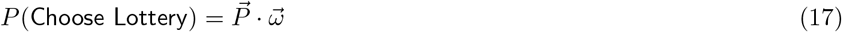

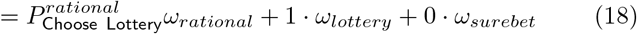

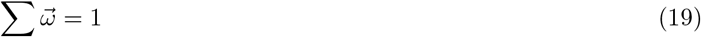

#### Model fitting

We estimated the posterior distribution over model parameters with weakly informative priors using the `rstan` package (v2.21.2; Stan Development Team, 2020). `rstan` is the R interface of Stan (Stan Development Team, 2020), a probabilistic programming language that implements a Hamiltonian Monte Carlo (HMC) algorithm for Bayesian inference. The prior over the utility exponent *ρ* was *Lognormal*(log(0.9), 0.4), a weakly informative prior that prefers *ρ* to be close to risk-neutral. The prior over noise *σ* was *Gamma*(6, 3). The prior over the mixing weights 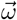 was a Dirichlet distribution with the concentration parameter *α* = [6, 2, 2]. The resulting *ω_rational_* distribution was broad and had the mean of 0.6, both *ω_lottery_* and *ω_surebet_* distribution had the mean of 0.2. By attributing more weight to the rational agent over the habitual agents, the prior reflected our selection of the experimental animals - only the ones with good psychometric curves were included. Four Markov chains with 1000 samples each were obtained for each model parameter after 1000 warm-up samples. The *R* convergence diagnostic for each parameter was close to 1, indicating the chains mixed well.

#### Inactivation mixture model

To estimate the effects of inactivations in our three-agent framework, we constructed a different version of the three-agent model, which considered three datasets from each rat simultaneously: bilateral FOF, bilateral PPC and a control dataset. The model’s raw parameters included *ρ_base_*, a parameter for *ρ* in the log space, its prior was *Lognormal*(log(0.9), 0.4); *σ_control_*, as *σ* in the original version, with a prior of *Gamma*(6, 3); *ω*_1_, with a prior of 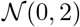, equivalent to *ω_rational_* after a *logistic* transformation; and *ω*_2_ with a prior of 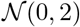, representing the proportion of the surebet agent in 1 – *logistic*(*ω*_1_) after the *logistic* transformation itself, where *logistic*(*x*) = 1/(1 + *e^-x^*):

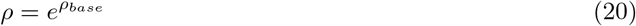

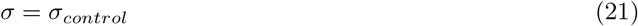

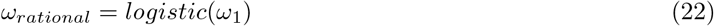

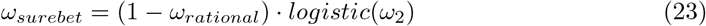

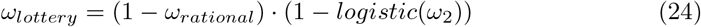

The re-implementation of the model was done so that the effects of silencing could be more easily interpreted. In particular, it was not clear to us how to implement a ‘zero-centered’ bias to the mixture parameter, *ω* as a Dirichlet. Also, for model estimation, the effects of inactivation should bias parameters in a space where support is guaranteed. For example, ensuring that *ρ* > 0; *σ* > 0;0 ≤ *ω* ≤ 1.

For each inactivation dataset, we added a new parameter for each raw parameter in order to estimate the effects of inactivation:

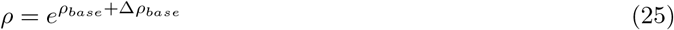

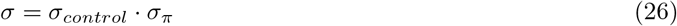

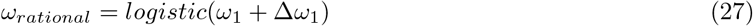

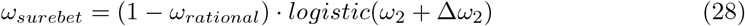

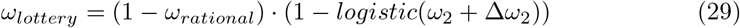

where Δ*ρ_base_* denotes the change in *ρ* in the log space, it had a prior of 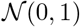; *σ_π_*, with a prior of *Lognormal*(0, 0.1), represents how the infusions could scale noise; Δ*ω*_1_ and Δ*ω*_2_ fit potential changes in *ω*_1_ and *ω*_2_ before the *logistic* transformation, respectively.

We constructed two other variants of the inactivation model for model comparison. For the *ρ*-only model, both Δ*ω*_1_ and Δ*ω*_2_ were fixed to be 0 during fitting. For the *ω*-only model, Δ*ρ_base_* was fixed to be 0.

#### Synthetic datasets

To test the validity of our model, we created synthetic datasets with parameters generated from the prior distributions described above. The three-agent model was fit to the synthetic datasets, and it was able to recover the generative parameters accurately (Figure S7A). This assured that our model can capture the behavior well and has no systematic bias in estimating the parameters.

#### Mixture model prediction confidence intervals

To generate model predictions in between the actual lottery lottery magnitudes (as in Figure 4B), we generated a synthetic dataset with narrowly-spaced lottery magnitudes (incremented by 1). Then, we sampled parameters from the estimated posteriors and computed the probability of choosing the lottery given the synthetic offers. The resulting output is a `n_iter` × `n_lott_mag` matrix, where `n_iter` is the number of Markov samples and `n_lott_mag` is the length of unique lottery magnitudes. Finally, binomial 80%, 90%, and 95% confidence intervals for each lottery offer were estimated by taking the 10% and 90%, 5% and 95%, and 2.5% and 97.5% percentile of `n_iter` predicted choices, respectively.

#### Mixture model comparison

To understand which model describes the inactivation results best, we performed 10-fold cross-validation of the MCMC fits of each model. For each fold, the model first fit on the training data, containing 90% of the original data from each condition (control and bilateral FOF). We then computed (in the `generated quantities` block) the log predictive densities by passing in the held-out data, using the posterior draws conditional on the training data. As the training and testing data are independent, the log predictive density coincides with the log likelihood of the test data. To evaluate the predictive performance, we computed the expected log pointwise predictive density (ELPD) using the test data (Vehtari et al., 2017). As the definition of ELPD incorporates the true generating process of prediction that is unknown, in practice, ELPD is approximated by computing the log predictive density using draws from the posterior samples

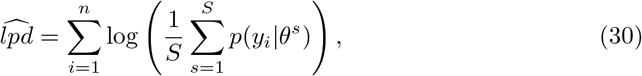

where *n* is the number of test trials, *θ^s^* is the s-th parameter sample from the posterior, and *p*(*y_i_*|*θ^s^*) is the log predictive density of the i-th test trial computed using the s-th parameter sample. Intuitively, the closer ELPD is to 0, the higher the model predictive accuracy.

### Dynamical model

We generated a 6-node rate model as a potential mechanism for understanding how muscimol inactivation of the FOF could cause a reduction in lottery choices via a change in the curvature of the utility function. The activity of the six nodes, *X*, are governed by the following equations, where *v* is the magnitude of the lottery and the *i* in *g*(*v,t,i*) represents the node index (1-6). Simulation was done using Euler’s method in Julia 1.6.0,^50^:

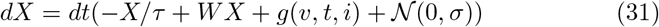

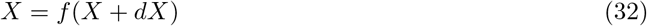

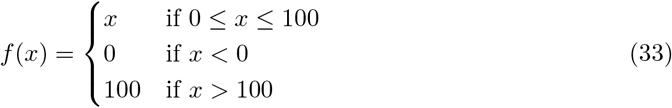

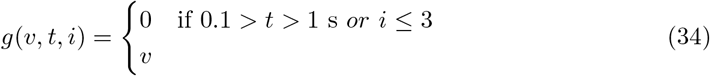

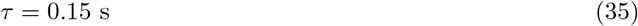

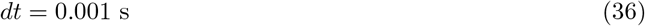

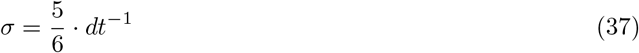

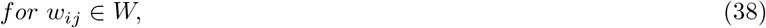

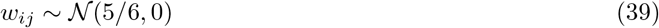

We began the simulation of each trial a few seconds before the input was turned on, to allow the network to reach its baseline fixed-point. We examined different instantiations of this model by generating the weight matrix, *W*, from different random seeds. Many (but not all) of these networks gave qualitatively similar results.

### Electrophysiology

Six rats were implanted with movable silicon probes (Cambridge NeuroTech) in either left or right FOF for single-unit electrophysiology data collection (four rats implanted contralateral to the lottery side, two rats implanted to the lottery side). The silicon probes were adhered to nano-drives (Cambridge NeuroTech) with super-glue. Following the same procedure described above, we marked the location of the FOF (+2.5 AP, ±1.4 ML mm from Bregma), and then a 1.5mm craniotomy was drilled, followed by an entire dura resection. The craniotomy was then filled with saline saturated Gelfoam to protect the brain tissue while the skull was coated with a thin layer of C&B Metabond (Parkell, Inc; New York) and a 1-3mm high moat built around the craniotomy using the Absolute Dentin (Parkell, Inc; New York). Then, the adjustable nano-Drive assembled silicon probes was mounted to the stereotax. Ground wires were soldered to titanium ground screws located above primary visual cortex. The silicon probe was slowly lower into the brain until all the recording sites were immersed into the tissue (1.3mm DV for the H3 probes, and 0.5mm DV for the E probes). The craniotomoy was filled with Dura-Gel (Cambridge NeuroTech), and then microdrive was cemented to the skull with Absolute Dentin.

After the surgery, the rats recovered for six days with *ad libitum* access to food and water. The recording sessions began at the 7th day post operative. Neural activity was digitized at 30 kHz, amplified and bandpass filtered at 0.6-7500Hz using a 64 channel intan headstage (RHD2164, Intan Technologies, https://intantech.com/files/Intan_RHD2164_datasheet.pdf), the SPI cable of the intan headstage was tethered to a commutator to allow free spinning(Shenzhen Moflon Technology, MMC250), and all the raw data was processed using the Open Ephys acquisition board (https://open-ephys.org/acquisition-system/eux9baf6a5s8tid06hk1mw5aafjdz1) connected to a computer to visualize and store the neural signals.

During the recording, a serial TTL messages encoding the current trial number was sent from our behavioral control hardware to to the acquisition system to synchronize the neural signal with the behavior data. The probes were turned down ~ 100μm every day for 4-6 days until the white matter was reached.

Offline spike sorting was performed by using **Kilosort v2** with the default setting. Spike clusters were manually curated using Phy. The quality metrics and waveform metrics for sorted units were computed using `ecephys spike sorting` https://github.com/AllenInstitute/ecephys_spike_sorting^51^. Specifically, we selected units with averaged firing rate >1Hz, SNR(signal to noise ratio) >1.5, and a presence ratio >0.95 over the course of recording sessions.

#### Data analysis

The spike times were aligned with the sound cue to plot the spike raster and peri-stimulus time histograms (PSTHs) in a 1.2 s time window (−0.1 s before the cue onset, and 0.1 s after the fixation end) with the bin size set to 10 ms resolution and smoothed with a causal half gaussian kernel with standard deviation of 20 ms.

To determine whether the firing of the cell could predict the upcoming choice selection during the fixation period, we counted the spikes for the late fixation period (0.5 - 1 s after the cue onset). We then ran a mixed-effects linear regression to see how task choice affected neural responses in each time window. Single cell mixed-effects linear models were fit in Matlab (Mathworks) using *fitlme* with the following formula:

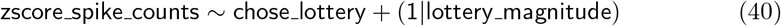

Where zscore_spike_counts was z-scored spike counts for that time window, chose_lottery was a binary data, which was 1 if the lottery was chosen on that trial, otherwise it was 0. lottery_magnitude were six relative reward values (0.5, 2, 4, 8, 16, 32) corresponding to the six distinct sound cues. A *p* < 0.05 for the coefficient of the fixed parameter chose_lottery was used to identify a choice selectivity cell.

Another mixed-effects linear regression was implemented to evaluate the contribution of different lottery magnitudes to spike firing in FOF regardless of chose_lottery, this was fit using Matlab function *fitlme* with the following formula:

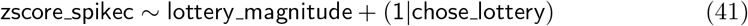

A *p* < 0.05 for the coefficient of the fixed parameter lottery_magnitude was used to identify a lottery tuning cell.

## Results

### Behavior

We trained rats on a ‘risky-choice’ task where they choose between a lottery and a surebet choice on each trial. The value of the lottery on each trial was indicated by an auditory cue (Figure 1A,B). In this paper, we only present behavior from sessions after the animal was implanted with cannulae for experiments. Unless otherwise specified, control sessions were the sessions from the day before the infusion sessions. The animals’ choices were largely consistent with a utility-maximizing strategy: they had relatively few violations of first-order stochastic dominance (i.e. they chose the surebet when the lottery magnitude was less than the surebet magnitude) and they increased the proportion of lottery choices monotonically with increasing expected value (Figure 1B-D). Visual inspection of the psychometric curves shows that six of the rats were risk-averse and two were close to risk-neutral (Figure 1D). On average, each rat completed 82 choices in a control session (Figure 1E).

### Effects of silencing FOF and PPC on the risky choice task

All animals experienced three different types of inactivations (left, right and bilateral) in two brain areas (FOF & PPC). In total, we include 7456 choice trials from 127 infusions sessions into the FOF and PPC of 8 rats. The details of region, order and dosage of the infusions for each rat are shown in Figure S2.

### FOF silencing shifted choices away from the lottery

Bilateral silencing of the FOF (Figure 2A) resulted in a dose-dependent reduction in lottery choices (Figure 2B; for individual subjects see Figure S3A). A generalized-linear mixed-effects model (GLMM) of the bilateral infusions found a significant main effect of muscimol dosage (*β_dose_* = −3.18 ± 0.92, *p* < 0.01). The mean indifference point (in units of *EV_lottery_* – *EV_surebet_* = *μL* of water) shifted from 50.92 ± 11.56 in control to 154.43 ± 23.49 under 0.3 *μ*g muscimol (T_8_ = −3.95, *p* < 0.001). In other words, inactivating bilateral FOF is equivalent to adding around 100 *μ*L to the surebet. There also was a significant decrease in slope (*βEV_lottery_-EV_surebet_:dose* = −0.08 ± 0.02, *p* < 0.001). Bilateral silencing of the FOF did not consistently change animal’s reaction time, defined as the time from center port withdrawal until a choice port poke (Linear mixed-effects model, LMM, *β_dose_* = 0.21 ± 0.29, *p* = 0.574). However, there was a significant slowing effect in three animals: 2152 (*β_dose_* = 2.33 ± 0.42, *p* < 0.001), 2153 (*β_dose_* = 1.31 ± 0.39, *p* < 0.01) and 2166 (*β_dose_* = 1.22 ± 0.25, *p* < 0.001), possibly due to muscimol spillover into the adjacent M1 area (Figure S2D). Overall, the slowing effect from bilateral FOF inactivation was less reliable across animals than the effect on choice (Figure S4A), suggesting the effect on choice was not primarily driven by changes in movement.

Unilateral infusions had a smaller effect compared to bilateral infusions (Figure 2C). Infusions of 0.3 *μg* muscimol into the left and right FOF resulted in small but significant decrease in the slope (*β_EV_right_–EV_left_:left_* = −0.01 ± 0.003, *p* < 0.001; *β_EV_right_–EV_left_:right_* = −0.01 ± 0.004, *p* = 0.049). These results were surprising for two reasons. First, we expected an ipsilateral bias, but both left and right infusions shifted animals slightly to choose leftward choices. As seven out of eight animals had the surebet port assigned on the left, it is possible that the decrease in choosing *right* after silencing either side of the FOF was, in fact, a partial effect of bilateral FOF inactivation (a decrease in choosing the lottery). Second, these effects were weak compared to the large ipsilateral biases caused by unilateral FOF silencing in previous tasks^41,52,53^. The discrepancy may be due to the memory component in previous tasks, whereas the risky-choice task does not have one. Overall, unilateral infusions in FOF did not change animal’s reaction time (LMM, *β_L_* = 0.07 ± 0.07, *p* = 0.44; *β_R_* = 0.01 ± 0.07, *p* = 0.89).

### PPC silencing had minimal effect on the risky choices

Bilateral silencing of PPC resulted in minimal effect on the risky-choice behavior (*βdose* = −0.60 ± 0.57, *p* = 0.29; Figure 2E, Figure S3C). To test for any lateralized effects from unilateral PPC infusions, we performed a second GLMM test where the choice on each trial was a logistic function of *EV_right_* – *EV_left_*, infusion side and their interaction as fixed effects. No significant effects were found on the group level (Figure 2F, all *p* > 0.5). To probe whether perturbation of FOF could reveal an effect of PPC inactivation, we inactivated unilateral FOF (0.3 *μ*g) while unilaterally inactivating PPC with 0.6 *μg* muscimol. The simultaneous inactivation, still, had no significant effect on the behavior (Figure 2D, all *p* > 0.5). Overall, the results suggest that PPC inactivation was ineffective in biasing risky choices. Thus, our hypothesis that the PPC may be involved in economic decisions because they are an expression of an internal preference was not supported.

### Bilateral PPC inactivation did not impair learning

The GLMM analysis above shows that the rat PPC was not causally involved in the risky choice task. However, numerous studies have found that neural activity in PPC correlates with decision variables in both perceptual and economic tasks. The question thus remains, what is the purpose of these decision-related signals in PPC? Recently, Zhong et al.^42^ found that PPC silencing impaired the ability of mice to re-categorize previously experienced stimuli based on a new category boundary in an auditory decision-making task. Moreover, after the stimuli were re-categorized, PPC activity was no longer required for performance. Motivated by their findings, we tested whether PPC was necessary for re-categorizing stimuli in our task. To do so, we employed a model-based change in the surebet magnitude that effectively shifted the decision boundary without changing the frequency-to-lottery mapping (Figure 3A). As such, some frequencies that had led to mostly lottery choices now led to mostly surebet choices (and vice-versa, depending on the direction of the shift). To estimate the required shifts, we first fit the three-agent model on data from the past 14 sessions. We then used the fit to generate synthetic choices on different surebet magnitudes, until we found the one that resulted in a shift in the overall probability of choosing lottery (*p*(Choose Lottery)) close to the target (drawn uniformly from *±U*(0.2, 0.3); see details in Methods). To familiarize animals with the new paradigm, their surebet magnitudes were changed weekly for two weeks. Two out of six animals failed to show appropriate adaptation of behavior following change in surebet magnitude; they were excluded from analysis in this section. The other four animals reliably shifted their choices more towards surebet when its magnitude increased, and more towards lottery when surebet magnitude decreased (See example animal in Figure 3B, all other animals in Figure S6)

After two weeks, on the day of surebet change, we infused 0.6 *μ*g muscimol into each side of PPC in these four animals before the task. The animals learned the new surebet magnitude and adjusted their behavior in both control and PPC inactivation sessions (see example animal in Figure 3C). To quantify the effectiveness of the surebet shift and the potential contribution of the PPC to the shift, we compared the predicted shifts to the actual shifts (Figure 3D). Bilateral PPC inactivation did not impair the learning of new surebet magnitudes. In fact, we found the opposite. Normally, the actual shifts were smaller than the predicted shifts (Figure 3D, blue). On days when the PPC was silenced the actual shift was closer to the predicted shift (*β_predicted_shift:PPC_* = 0.251 ± 0.091, *p* = 0.011; Figure 3D, yellow). Thus, our results do not support the hypothesis that the PPC is required for shifting category boundaries: i.e. categorizing a lottery as being better or worse than the surebet. Instead, the smaller than expected shifts (Figure 3d-blue dots) could be explained as a contraction bias: the large shifts were unexpected given subjects’ prior beliefs about the magnitude of the surebet. Silencing the PPC (Figure 3D-yellow dots) reduced this contraction bias; i.e. weakened the influence of the prior on the behavior^54^.

### Unilateral PPC inactivation biases ‘free’ choices

In order to establish that our infusions into PPC were effective, after completing all of the experiments reported above, we added a ‘free’ trial type as in^41^. On a free trial, both the surebet port and lottery port were illuminated with blue-LEDs after fixation, accompanied by a brief neutral tone. The animals were rewarded twice the magnitude of the surebet reward regardless of which port they chose (Figure 3E). These types of trials have been demonstrated to be sensitive to unilateral silencing of the PPC^41,55^. We randomly intermixed 11% free trials with 22% forced trials and 67% choice trials on the control days. After a few sessions with the new trial type, rats expressed a consistent bias on the free trials and still performed the choice trials in a utility-maximizing way. The proportion of free trials was increased to 50% on the infusion day, with the rest being 12.5% forced trials and 37.5% choice trials. Infusions of muscimol (0.6 *μ*g) into one hemifield of PPC (opposite to the animal’s preferred side) produced a substantial ipsilateral bias on free trials (Figure 3F; *β_infusion_* = 1.19 ± 0.50, *p* < 0.05). The ipsilateral bias in free trials was observed even while, consistent with our previous PPC inactivation results, there was no ipsilateral bias on the interleaved choice trials (Figure 3G,H; *β_infusion_* = 0.18 ± 0.14, *p* = 0.189). These free trial inactivation results provided a clear positive control for our PPC inactivations, demonstrating that the lack of effect on choice trials was not caused by a technical issue (like clogged cannula).

### A three-agent mixture model of risky choice

While the effects of FOF silencing confirmed its role in decisions under risk (Figure 2B), the GLMM results did not provide insight into the specific role that the FOF might play. To better understand animal behavior in the task and the role of the FOF, we developed a three-agent mixture model (Figure 4A). The first agent was a ‘rational’, utility-maximizing agent ^37^ with two parameters: *ρ*, which controls the shape of the utility function (*U* = *V^ρ^*); *σ*, which captures the decision noise. The other two agents were stimulus-independent agents which either habitually chose the lottery or the surebet. The relative influence of the agents is controlled by their mixing weights *ω*, where 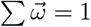. The choice on each trial is thus a weighted outcome of the ‘votes’ of three agents, each implementing a different strategy. We estimated the joint posterior over the parameters for each subject separately using Hamiltonian Monte Carlo sampling in Stan^56^ and validated that the model can correctly recover generative parameters from synthetic data (Figure S7A). Details of the modeling, including the priors, can be found in the Methods section. The motivation for developing the mixture model was that the animals’ choices, while clearly sensitive to the lottery offer, showed some stimulus-independent biases. In other words, even for the best lottery they sometimes chose the surebet and for the worst lottery (which had a value of 0) they sometimes chose the lottery. For example, subject 2156 has a psychometric curve that asymptotes in a way that is inconsistent with a pure utility-maximizing strategy (Figure 4B).

Trial history effects could have been incorporated by allowing model parameters to vary depending on the outcome of the previous trial as in^26^. However, our animals seemed to understand that the lottery offer was independent across trials, and we did not see any statistically significant effects of previous trial’s outcome on choice in control sessions (GLMM, *β_lottery-win_* = 0.20 ± 0.12, *p* = 0.08;

*β_lottery-lose_* =0.17 ± 0.09, *p* = 0.08). For this reason, we decided to formulate the three-agent mixture model without trial history parameters. Our animals’ behavior stood in contrast to a substantial number of published results demonstrating strong trial history effects in rodent decision-making even when the optimal strategy is to only use information on the current trial e.g.^26,57,58^. We speculate that an important difference is that in traditional rodent two-alternative forced-choice tasks, the rewards were delivered at the choice ports, but in our task all rewards were delivered at a single reward port but for counter examples where there is history dependence despite using a single reward port, see^59,60^.

The three-agent model fit the control behavior well (see two example animals in Figure 4B, all animals in Figure S7B). For example, rat 2156 chose the lottery about 10% of the time for the 4 worst lotteries. This behavior is not well described by a ‘pure’ utility-maximizing model. All animals had a decelerating utility function (95% C.I. of *ρ* < 1 for all animals; Figure 4C). Note that the *effective* risk-preference is influenced by both *ρ* and *ω*. For example, the indifference point of 2152 is close to 0, implying that it is effectively risk-neutral (Figure 4B). However, this comes from its bias towards choosing the lottery (*ω_lottery_* = 0.16) balancing its decelerating utility function (*ρ* = 0.62; Table 1). The animals had small but varying levels of decision noise (*σ* = 1.00 [0.35 3.40], median and 95% C.I. of concatenated posteriors across animals), indicating that they were sensitive to water rewards just a few *μ*L apart. Their choices were guided mostly by the rational agent (*ω_rational_* = 0.82 [0.65 0.95]), with little influence from the lottery agent (*ω_lottery_* = 0.10 [0.01 0.31]) and the surebet agent (*ω_surebet_* = 0.05 [0.01 0.23]).

**Table 1.** Fits from the three-agent inactivation model. Statistics were computed using the parameter posteriors from the three-agent model fit to the control, 0.3 *μ*g per side bilateral FOF inactivation, and 0.3 *μ*g per side bilateral PPC inactivation dataset simultaneously. The median of the parameter posterior distribution is reported along with its 95% confidence interval in brackets.

| | | $\rho$ | $\sigma$ | $\omega_{rational}$ | $\omega_{lottery}$ | $\omega_{surebet}$ |
| --- | --- | --- | --- | --- | --- | --- |
| **2152** | Control | 0.62 [0.59, 2.07] | 1.61 [1.16, 3.06] | 0.82 [0.64, 0.87] | 0.16 [0.11, 0.18] | 0.02 [0.01, 0.20] |
|  | FOF | 0.38 [0.11, 9.07] | 1.54 [1.10, 3.21] | 0.97 [0.14, 1.00] | 0.02 [0.00, 0.28] | 0.00 [0.00, 0.70] |
|  | PPC | 0.66 [0.58, 0.78] | 1.62 [1.12, 3.12] | 0.78 [0.64, 0.87] | 0.14 [0.06, 0.21] | 0.09 [0.04, 0.20] |
| **2153** | Control | 0.67 [0.65, 2.41] | 2.52 [1.39, 4.63] | 0.79 [0.62, 0.86] | 0.20 [0.14, 0.23] | 0.00 [0.00, 0.18] |
|  | FOF | 0.42 [0.13, 6.71] | 2.32 [1.38, 4.77] | 0.97 [0.03, 1.00] | 0.02 [0.00, 0.22] | 0.00 [0.00, 0.90] |
|  | PPC | 0.62 [0.56, 0.71] | 2.37 [1.36, 4.79] | 0.89 [0.80, 0.99] | 0.09 [0.01, 0.16] | 0.02 [0.00, 0.06] |
| **2154** | Control | 0.45 [0.42, 0.50] | 1.05 [0.73, 1.21] | 0.96 [0.82, 0.98] | 0.02 [0.01, 0.06] | 0.03 [0.00, 0.14] |
|  | FOF | 0.35 [0.28, 0.65] | 0.84 [0.69, 1.27] | 0.99 [0.46, 1.00] | 0.01 [0.00, 0.12] | 0.00 [0.00, 0.48] |
|  | PPC | 0.43 [0.39, 0.52] | 0.90 [0.68, 1.27] | 0.93 [0.72, 0.97] | 0.07 [0.02, 0.15] | 0.00 [0.00, 0.19] |
| **2155** | Control | 0.39 [0.36, 0.42] | 0.48 [0.43, 0.65] | 0.84 [0.77, 0.96] | 0.05 [0.03, 0.07] | 0.10 [0.00, 0.18] |
|  | FOF | 0.33 [0.31, 0.42] | 0.49 [0.41, 0.66] | 0.96 [0.68, 1.00] | 0.02 [0.00, 0.04] | 0.01 [0.00, 0.31] |
|  | PPC | no data | no data | no data | no data | no data |
| **2156** | Control | 0.55 [0.46, 0.59] | 0.19 [0.22, 0.56] | 0.71 [0.66, 0.76] | 0.14 [0.12, 0.17] | 0.15 [0.10, 0.19] |
|  | FOF | 0.22 [0.05, 2.34] | 0.15 [0.21, 0.58] | 0.56 [0.05, 0.99] | 0.29 [0.00, 0.51] | 0.15 [0.00, 0.81] |
|  | PPC | 0.50 [0.42, 0.66] | 0.21 [0.22, 0.59] | 0.61 [0.56, 0.86] | 0.09 [0.03, 0.12] | 0.30 [0.08, 0.36] |
| **2160** | Control | 0.66 [0.56, 0.67] | 0.59 [0.48, 1.17] | 0.76 [0.72, 0.82] | 0.21 [0.16, 0.24] | 0.03 [0.01, 0.06] |
|  | FOF | 0.32 [0.33, 0.66] | 0.58 [0.47, 1.15] | 0.98 [0.53, 1.00] | 0.02 [0.00, 0.07] | 0.01 [0.00, 0.43] |
|  | PPC | 0.54 [0.50, 0.70] | 0.50 [0.46, 1.22] | 0.82 [0.75, 0.96] | 0.17 [0.03, 0.18] | 0.01 [0.00, 0.12] |
| **2165** | Control | 0.45 [0.43, 0.50] | 0.34 [0.41, 0.92] | 0.88 [0.84, 0.97] | 0.07 [0.01, 0.06] | 0.06 [0.01, 0.12] |
|  | FOF | 0.29 [0.11, 0.80] | 0.37 [0.38, 0.94] | 0.99 [0.10, 1.00] | 0.01 [0.00, 0.08] | 0.00 [0.00, 0.87] |
|  | PPC | 0.43 [0.37, 0.53] | 0.29 [0.41, 0.96] | 0.98 [0.82, 1.00] | 0.01 [0.00, 0.06] | 0.01 [0.00, 0.15] |
| **2166** | Control | 0.43 [0.43, 0.54] | 0.61 [0.51, 1.51] | 0.92 [0.87, 0.97] | 0.07 [0.02, 0.10] | 0.01 [0.00, 0.05] |
|  | FOF | 0.22 [0.14, 1.01] | 0.49 [0.47, 1.55] | 1.00 [0.21, 1.00] | 0.00 [0.00, 0.10] | 0.00 [0.00, 0.75] |
|  | PPC | 0.57 [0.28, 0.88] | 0.68 [0.50, 1.58] | 0.82 [0.51, 0.99] | 0.03 [0.01, 0.37] | 0.15 [0.00, 0.23] |

### Bilateral FOF inactivation reduced the utility exponent *ρ*

In order to quantify how the infusions influenced model parameters, we constructed a new version of the three-agent model that fit the 0.3 *μg* bilateral FOF and 0.3 *μg* bilateral PPC infusion data as perturbations of the control dataset for each subject (Figure 5A & Figure S9). We chose priors for the effects of perturbation such that the model favored no effect of inactivation (i.e. zero mean for shifts and one mean for scaling effects). Bilateral PPC infusion led to no reliable changes across subjects for all parameters, which was consistent with the results from the GLMM (Figure S9 & Table 1). From the GLMM (and visual inspection), we knew that bilateral FOF silencing substantially shifted the subjects to being effectively more risk-averse. Indeed, the model-based analysis showed that overall animals had a reduction in *ρ* compared to the control fits (Figure 5B-bottom). However, there was also a fairly reliable decrease in the contribution of the lottery agent (Figure 5B-top).

**Figure 5.**
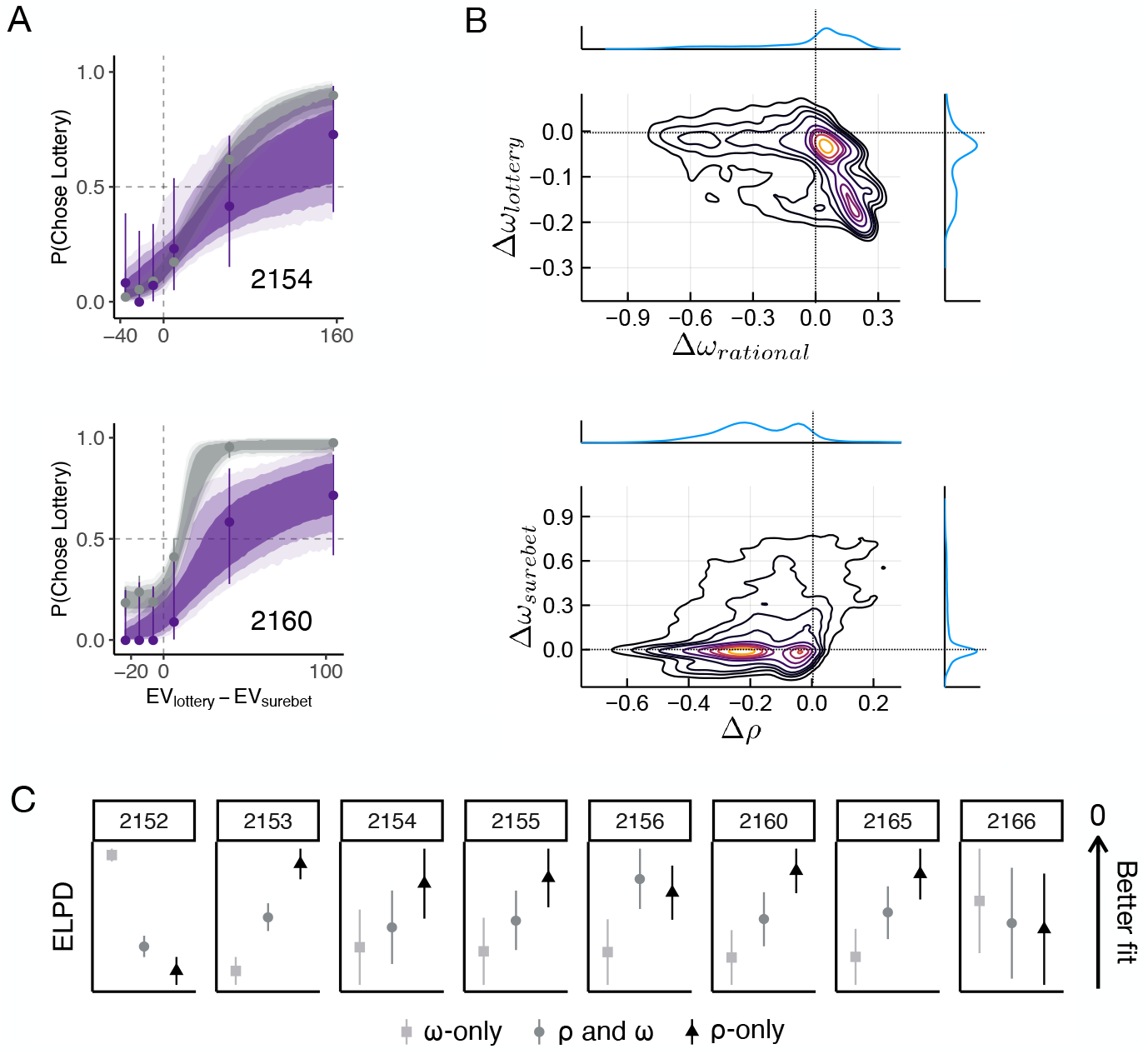
Bilateral FOF inactivation reduced the exponent of the utility function. **A.** Psychometric curves for two example animals. The circles with error bars are the binned mean and 95% binomial confidence intervals. The ribbons are model predictions generated using the fitted parameters (control–gray, bilateral FOF inactivation–purple). The dark, medium and light shade represent 80%, 95% and 99% confidence intervals, respectively. Similar plots for all animals are shown in Figure S8. **B.** Contour plot of posterior samples of the deviations of the model parameters under bilateral FOF inactivation from control. The most reliable findings across animals were decreases in *ρ* and *ω_lattery_*: *P*(Δ*ρ* < 0) = 0.856; *P*(Δ*ω_lottery_* < 0) = 0.844. Individual fits are shown in Figure S9. The lottery bias was small in the control sessions, so a decrease in that bias cannot account for the large shift towards choosing the surebet during bilateral FOF inactivation. **C.** Ten-fold cross-validation results comparing three model variants: where *ρ* but not *ρ* was allowed to change under the inactivation dataset, both *ρ* and *ω* were allowed to change, and *ρ* but not *ω* was allowed to change. The points with error bars are the expected log posterior density (ELPD) and its standard error on each animal’s dataset. The *ρ*-only model was preferred to the *ω*-only model in 6 out of 8 animals.

To test whether the reduction in lottery choices was actually due to a decreased *ρ* rather than changes in *ω*, we constructed two variants of the inactivation model and compared them using 10-fold cross validation (see Methods for details). The ‘ρ-only’ model had parameters allowing *ρ* and σ to shift, but not any *ω* parameters to change under inactivation. Similarly, the ‘ω-only’ model had parameters allowing only *ω* and σ but not *ρ* to change. The ‘ρ and *ω*’ model was the standard inactivation model that allowed every parameter to change under inactivation. We found that in 6 out of 8 subjects, model comparison result preferred the *ρ*-only model over the *ω*-only model (Figure 5C). The *ω*-only model was strongly preferred only in one subject’s dataset (2152). Taken together, these results suggest that the most parsimonious interpretation of the inactivation-induced effect is a reduction in the utility exponent, *ρ*.

How could silencing the FOF change the exponent of the utility function? Previous silencing and modeling results suggested that the FOF is part (1/6) of a distributed circuit for maintaining a prospective memory of choice^53^. Inspired by that finding, we constructed a 6-node rate model of a distributed circuit for encoding action-value, where the FOF represented one node in that network Figure 6A;^61^. Three nodes other than the FOF node received input representing the magnitude of the lottery. The all-to-all weight matrix was generated randomly, but the distribution of the weights was chosen such that the response of the network to the inputs was in the dynamic regime of the nodes (0 < *Hz* < 100). Other network parameters (noise *σ* and time-constant *τ*) were chosen to generate a control network response with reasonable dynamics (Figure 6B) that encoded the lottery value in the population activity of the network (Figure 6C, gray circles). In this regime, we found that silencing the FOF node scaled down the network’s responses. Note, that the scaling is not a trivial 1/6 reduction in the average firing rate but reflects the contribution of the FOF node to the overall network dynamics (Figure 6C-firing drops from 60 to 25 Hz for the largest lottery). We can think of this network as encoding the expected utility of choosing the lottery by transforming the lottery sound into *utils* (encoded as spike rate). At the time of the go-cue, this activity could become bistable: where the utility of the surebet determines the unstable fixed point similar to^62^. Alternatively, a downstream region could compare the output of this network with the remembered surebet utility. In any case, scaling down the input-output transform of the network (Figure 6C, purple circles) would shift the indifference point (the lottery that had the same activity level as the surebet comparator), which would, behaviorally, appear as a change in the power-law utility function *U* = *V^ρ^*. For the control network, the network approximates a function with *ρ* ≈ 0.76. After silencing the FOF node, the exponent of the utility functions shifted down, *ρ* ≈ 0.6 (Figure 6C). This dynamical model provides a mechanistic explanation for our finding that silencing the FOF with muscimol caused animals to avoid choosing the lottery (Figure 2B) through a change in the exponent of the utility function (Figure 5B).

**Figure 6.**
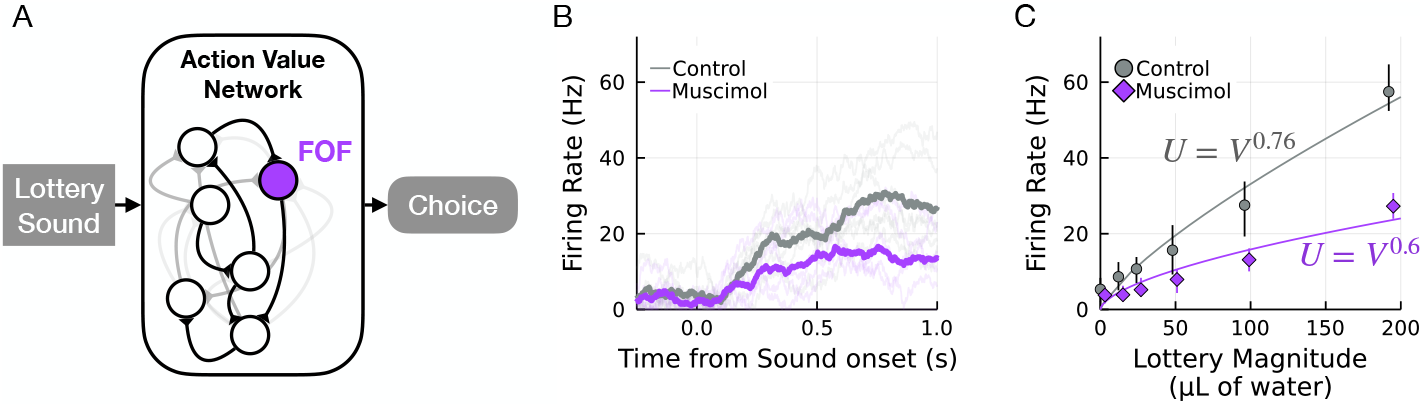
Dynamical model of FOF silencing. **A.** We implemented a 6-node rate model of a distributed action-value network with random connectivity 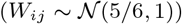. The FOF was 1 of the 6 nodes (in purple). The input to the network was the lottery magnitude. For the following plots the random seed for generating *W* was set to 131 and then that *W* was used for all further simulation, but similar results can be obtained with other W generated with the same statistics from a different seed. **B.** Example of the network response to lottery sound with magnitude of 96 *μL* under control conditions (with all the nodes active, in grey) and under FOF silencing (the FOF node is set to zero, in purple). The dark traces represent the mean network activity and the light traces represent the activity of the 6 individual nodes. **C.** Silencing FOF scales down the representation of the action-value of the lottery, which could explain the shift in *ρ*. We ran the network for 20 ‘trials’ of each lottery ∈ [0, 12, 24, 48, 96, 192] *μ*L. The grey circle are the mean and 95% CI for the network response in the control conditions and the purple diamonds are the mean and 95% CI for the network response when the FOF node is silenced. Fitting a power-law utility function, *U* = *V^ρ^* to the network activity gives *ρ* ≈ 0.76 for control, and after FOF silencing *ρ* ≈ 0.6. The thin lines are power-law utility functions that approximate the transformation from units of reward (*μ*L) to *utils* in spikes / second.

### Physiological evidence of a value map in FOF

Our dynamical model (Figure 6) suggested that neurons in the FOF should monotonically increase their firing rate with increased lottery values. To test this, we recorded single-unit activity from the FOF of six rats performing the risky choice task (Figure S10). We found many neurons whose activity was consistent with a value encoding in the service of decision-making (Figure 7A,B) as predicted by our muscimol results and our dynamical model. Specifically, during the fixation period, these neurons fired more on trials with higher lottery values, even when controlling for the choice of the animal (Figure 7B,C). Many neurons also fired more for lottery choices compared to surebet choices, even when controlling for lottery magnitude (Figure 7A). Out of 1690 neurons recorded, 423 (25.0%) significantly correlated with ΔEV, (out of these 63.6% were positively correlated, the rest were negatively correlated) even when controlling for choice. We also found that activity during fixation of 702 (41.5%) of neurons predicted the upcoming choice of the animal (controlling for ΔEV). 309 neurons (18.3%) significantly encoded both ΔEV and choice. Together with our muscimol, Bayesian modeling, and dynamical model, this data strongly supports the view that the activity in the FOF represents action-values during planning as well as actions *per se*.

**Figure 7.**
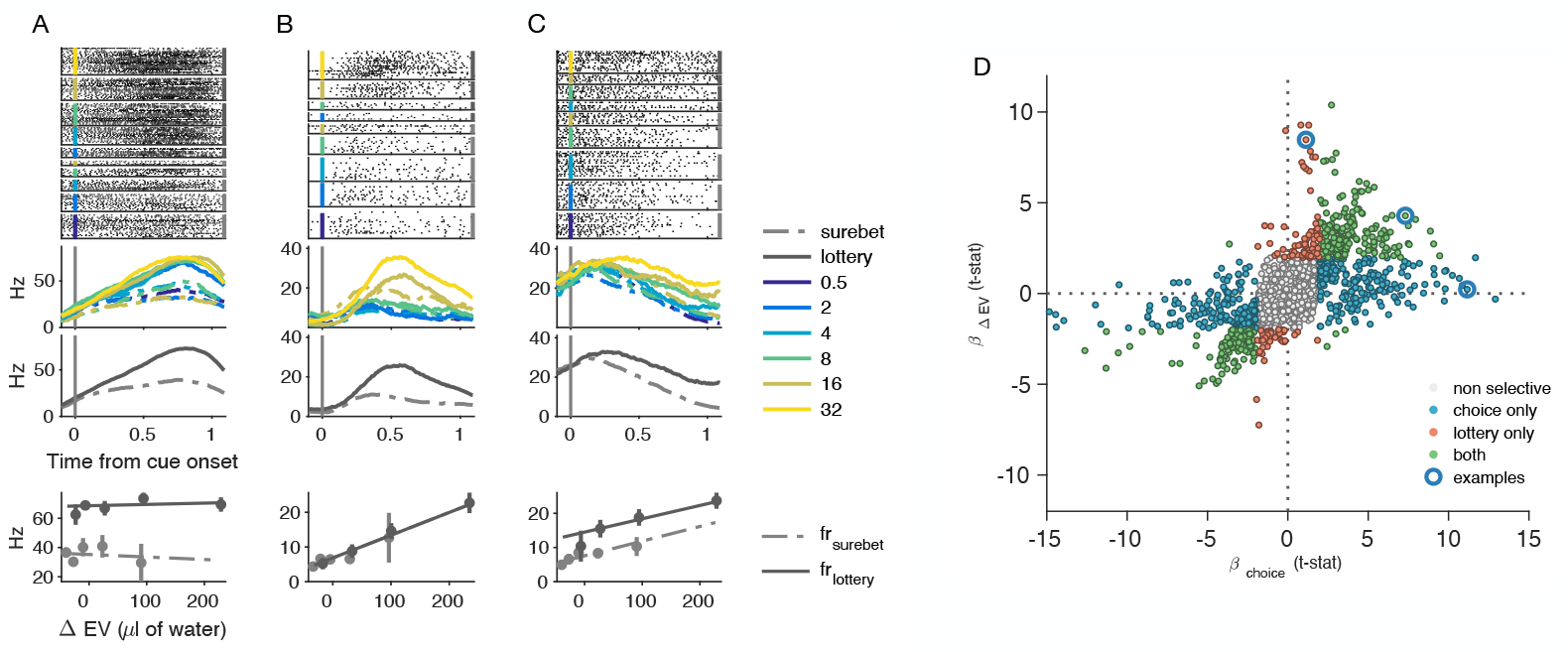
Neural activity in FOF encoded lottery value and upcoming choice action during the fixation period. **A-C** Example neurons. Spike raster plots and peri-stimulus time histograms (PSTHs) of three representative neurons shows heterogeneous dynamics during the fixation period. The rasters and PSTHs were aligned to the lottery cue onset, and were sorted by different lottery cues (indicated by different color) and upcoming choices (solid line for lottery choices, dashed line for surebet choices). The lower PSTH is sorted only on the basis of upcoming choices. The bottom row (Firing Rate vs. ΔEV) summarize the relationship between the neural activity, lottery value and choice. The lines are fits of a linear model (Hz Δ*EV*\*choice) to the data (dots with error bars representing standard error of the mean) **D.** Distribution of coefficients of the choice against ΔEV from the mixed effects linear models to characteristic the individual neuronal responses to upcoming choices or lottery values. Of the total 1690 recorded neurons, 23.5% (n=393) were tuned for upcoming choice alone, 6.8% (n=114) were purely tuned for the lottery values, while 18.3% (n=309) were tuned for both. Blue circled data showed where the three example neurons are located in the scatter plot.

## Discussion

The neurobiology of decision-making under risk has been studied extensively in humans, non-human primates and rodents. However, there has been a gap in task design between the human-primate and rodent experiments, that most rodent studies focused on unexpected uncertainty where choices often reflected their sensitivity of reward history rather than risk attitudes. Here, we developed a risky choice task for rats, where animals made cue-guided decisions between a lottery and a surebet option on a trial-by-trial basis under expected uncertainty, as in human and primate experiments but see^27,63^ for recent examples of rodent work on expected uncertainty. We developed the three-agent mixture model to decompose different elements of risk-preference, including *ρ* as exponent on the utility curve and 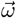 as the weights for the rational, lottery and surebet agents. Modeling results showed that all subjects had decelerating, i.e. risk-averse, utility functions and their decisions were influenced mostly by the rational agent. We tested how inactivations of two cortical regions, the FOF and PPC, influenced choices in our task. These regions have been studied extensively in perceptual decision-making, but this is the first test of their casual role in economic decision-making. Bilateral FOF inactivations produced a profound bias towards the surebet, while bilateral PPC inactivations had minimal effect on the behavior. Model-based analyses of the results indicated that, without the FOF, subjects had utility functions that were substantially shifted towards risk-aversion. We constructed a dynamical model to show that inactivating the FOF node can produce outputs similar to the effects observed, providing a mechanistic explanation. To further test the hypothesis that FOF neurons encode an action-value signal when rats perform the risky choice task, we recorded single neuron activity using silicon probes. Neurophysiological results revealed that among the recorded neurons in FOF, during the late fixation period, there were significant numbers of neurons reflected the upcoming choices, lottery values and both choice and lottery value signals. This fact, along with the muscimol inhibition experiments, suggested that FOF plays a critial role in encoding the action value to guide risky choice behavior. Finally, we found that PPC was not causally involved in the learning of new categorization boundaries, but may play a role in representing a prior about the value of the surebet.

### Role of FOF

Results from bilateral FOF inactivations show that FOF is an essential part of the circuitry underlying risky decision-making. Model-based analyses suggest that the change in behavior was likely due to a decrease in *ρ*, the curvature of the utility function, *U* = *V^ρ^*. However, there are many functional forms of decision-making under uncertainty that we did not test^32,45,64^, which could lead to different interpretations. Nevertheless, the change in *ρ* is consistent with the finding that inactivation of monkey supplemental eye field reduced risky choices and the change was characterized by a decreased utility exponent ^32^.

Using a dynamical model, we demonstrated that a shift in *ρ* can be caused by a partial inactivation of an action-value network whose activity guides choice^17^. This is similar to the theory that the FOF is part of a network for planning upcoming choice ^53^. In fact, the interpretation of FOF activity encoding action-value is consistent with the previous interpretation (planning movement), since in perceptual tasks, only the correct side is rewarded, making it difficult to disentangle action-value from movement-planning. This action-value network may be the locus of transformation from value to utility, with the network properties (e.g. whether the gain of the network is greater or less than 1) determining whether animals are risk-seeking or risk-averse. One key difference between our current and previous findings, is that previously, sensory-guided choices (i.e. trials with no working-memory requirement) were not affected by silencing FOF^41,53^. In Erlich et al.^41^, we posited that the FOF may be a bottleneck through which long-timescale integration of information could influence orienting decisions. Our results here suggest that the FOF may play a similar role for decisions that require integration of multiple attributes – in this case, lottery value and probability.

The idea that the FOF is part of an action-value network is largely consistent with the view that the FOF contributes to sensory-to-motor transformation^65–69^, but reinterprets those findings as sensory-to-value transformations. It is important to note that, in our task, the surebet value is stable across trials and only the lottery needs to be evaluated on a trial-by-trial basis. We predict that in a task where the lottery is stable across trials and the surebet value varies trial-by-trial (and is indicated by a cue), silencing the FOF would shift animals away from selecting the surebet, whose value would require transformation on each trial. In tasks that require transformations for both surebet and lottery e.g. ^63^, bilateral FOF silencing might result in increased decision noise.

Both left and right unilateral FOF inactivations led to a small bias towards leftward choices (Figure 2C), rather than an contralateral impairment, as previously reported^28,41,53^. However, in those studies, trials that did not require short-term memory were not biased by unilateral FOF inactivations, so the small effect is not particularly surprising. As 7 out of 8 animals had the surebet port on the left, the leftward bias can be interpreted as a weak bias towards choosing the surebet; i.e. a partial effect consistent with our bilateral silencing results.

To probe our hypothesis that FOF is an important part of action-value network, we analyzed neural activities in FOF from rats performing the risky choice task. Mixed-effects linear model showed strong encoding the upcoming choices during the late fixation period regardless of the sound cue for current lottery value. This result was consistent with our previous studies, which reported FOF encoding of planning for upcoming choices in FOF when rats performing a cue guided orienting task^52,53^. Our present studies also identified neurons that encoded lottery value regardless of upcoming choices. Moreover, 73.1% of these lottery-related neurons also encoded upcoming choice. Previous studies showed that value related neural signals have been found in the orbital frontal cortex and medial frontal cortex of non-human primates^70^, but no upcoming choice signal was found in these brain areas. In contrast, action values modulated neuronal activity has been reported in rat secondary motor cortex^71^, but not in the context of expected uncertainty. The majority of neurons were positively related to the lottery values, however, there were some value signals negatively related to the lottery values. While these ‘negative’ value neurons are not directly predicted by our model, it is quite common to observe neurons with opposite tuning^28,52,62^ while still having consistent effects of perturbations. Future experiments could examine whether the tuning is related to input or output projection patters^72^. Our neurophysiological results provide evidence for the first time that FOF encodes action value signals in the risky choice task. Our findings suggest that the FOF may be potential hub for the transformation of decisions in a ‘goods’ reference frame to an ‘action’ reference frame ^8^. Further analysis and optogenetic perturbation will be required to test how the value signal transformed into action in the FOF, and time course of the action value encoding and the related brain circuit.

### Role of PPC in risky choice

Activity in PPC has long been be associated with decision variables in economic choices. Platt and Glimcher ^29^ first showed that activity in monkey lateral intraparietal cortex (LIP), a visuomotor area within PPC, is sensitive to expected reward magnitude and probability. Subsequently, Dorris and Glimcher ^30^ found that neurons in monkey LIP encode relative subjective desirability of actions in a mixed-strategy game. Activity in human PPC also correlates with subjects’ risk preferences^73^. To date, we are not aware of any studies of the role of rodent PPC in economic decisions. Nonetheless, PPC encodes task-related variables during perceptual decisions e.g.^28,43,74^. We were frankly disappointed that neither unilateral nor bilateral inactivation had any effect on the risky choices. As far as we are aware, this is the first experiment that directly tests the causal role of PPC in economic choices.

The null result is reminiscent of the null effects of PPC inactivation in the Poisson clicks task in rats^41^, and of LIP inactivation in the random dot task in monkeys ^55^. The null effect was unlikely the result of insufficient inactivation, as unilateral PPC infusions led to a significant ipsilateral bias in the free choice trials, where the decisions were guided by internal side preference rather than action value (Figure 3E) and also significantly reduced the contraction bias in the surebet shifting experiment (Figure 3A). The free choice result replicates previous findings^41,55^, and is consistent with the literature on the role of rodent PPC in neglect^75,76^, providing a clear positive control for the inactivation experiments. Taken together, our results demonstrate that PPC is not strictly necessary for making utility maximizing choices under risk.

It has been argued that rodent PPC is important for visually-guided decisions but not other modalities. For example, pharmacological inactivation of PPC has been shown to impair mice’ performance in a visually-guided navigation task with a memory component ^77^, and in a multi-sensory perceptual task but when only using visual but not auditory cues ^43^. It was suspected that due to the anatomical proximity between PPC and the visual areas, these inactivation results may be caused by a muscimol spillover into the adjacent visual cortex. However, recent experiments utilizing optogenetics have shown that, targeted inactivation of PPC during the stimulus period disrupted performance only on the visual but not auditory trials ^78^, and impaired decision sensitivity in a visually-guided task with variable delays in mice^74^. As such, we cannot exclude the possibility that the null effect on risky choice may be due to the modality of stimuli used. However, silencing mouse PPC was shown to impair re-categorization of sounds in an auditory task^42^, so the controversy over the modality-specific role of PPC is not fully resolved.

### Role of PPC in learning

We have shown that inactivating PPC did not impair the animal’s ability to shift their choices in response to changes in the value of the surebet, inconsistent with the findings from Zhong et al. ^42^. There are some key differences in the design between our and their experiments that may explain this. First, their experiments were performed on head-fixed mice, whereas our rats were freely moving in the training box. Second, their mice had to categorize (or re-categorize) stimuli for the first time while PPC was inactivated. In contrast, the animals in our experiment were accustomed to changing surebet values for two weeks prior to PPC inactivation, understanding that the surebet value may change unexpectedly. Bucci and Chess ^79^ found that PPC-lesioned rats had trouble learning the association between light and food if previously the light was presented without food. Interestingly, normal learning was observed in another cohort of PPC-lesioned rats that were not pre-exposed to the light. They attributed the impairment to PPC’s role in directing attention to the stimulus whose meaning surprisingly changes. If it is the case that PPC activity is required for the learning of ‘surprising’ shifts in existing associations, the discrepancy between our experiment and Zhong et al.’s can be then resolved.

### Conclusion

Studies of neurobiology of economic choice in rodents have mostly focused on the reward-valuation circuit: including the amygdala^14,16^, basal ganglia^18^ and orbital-frontal cortex^21,27,63,80–82^. Here, we examined the casual contribution of two cortical areas associated with planning orienting decisions, the FOF and PPC, whose analogous primate regions have been implicated in economic decision-making^29,32^. We found that FOF is a critical node in the circuit for decisions under risk, while PPC is not. Our results show that FOF neurons represent expected value, even when controlling for choice and that it is causally involved in decisions under risk.

## Code and data sharing

Data and code required to replicate the results presented here are available at https://github.com/erlichlab/risk-fof-ppc-muscimol

## Author Contributions

J.M-M., S.D., and J.C.E. designed the risky-choice task and J.M-M. programmed the task and training pipeline, which was later amended by X.Z. for the experiments described in this manuscript. J.M-M. and J.C.E. initially devised the three-agent Bayesian model for the risky-choice task and collaborated with X.Z. in iterative implementation and testing. X.Z. did further independent work on the Bayesian model: performed the synthetic model checks, fit the model to the data, and generated the figures. X.Z. and J.C.E. adapted the Bayesian model to analyze the effects of infusions. X.Z. and J.C.E. designed the infusion experiments. Infusions were performed by X.Z. and C.B. and analyzed by X.Z. J.C.E generated the dynamical model and related figure. The electrophysiological experiments were designed by C.B. and J.C.E. Electrophysiological data was collected and analyzed by C.B. and J.L. The manuscript was written by X.Z., C.B. and J.C.E. with comments from the other authors.

## Acknowledgments

We thank Mingming Chen, Yidi Chen, Anyu Fang, NengNeng Gao, Yingkun Li and Cequn Wang for technical assistance related to building and maintaining lab infrastructure as well as training animals and assisting with infusions. We thank Liujunli Li for assistance with collecting electrophysiological data and for helpful comments on the manuscript. JCE acknowledges the support of the 111 project (Base B16018), the National Natural Science Foundation of China (NSFC), the support of the NYU-ECNU Institute of Brain and Cognitive Science at NYU Shanghai and the support of the funders of the Sainsbury Wellcome Centre: The Wellcome Trust and the Gatsby Charitable Foundatoin.

---

**Figure S1.**
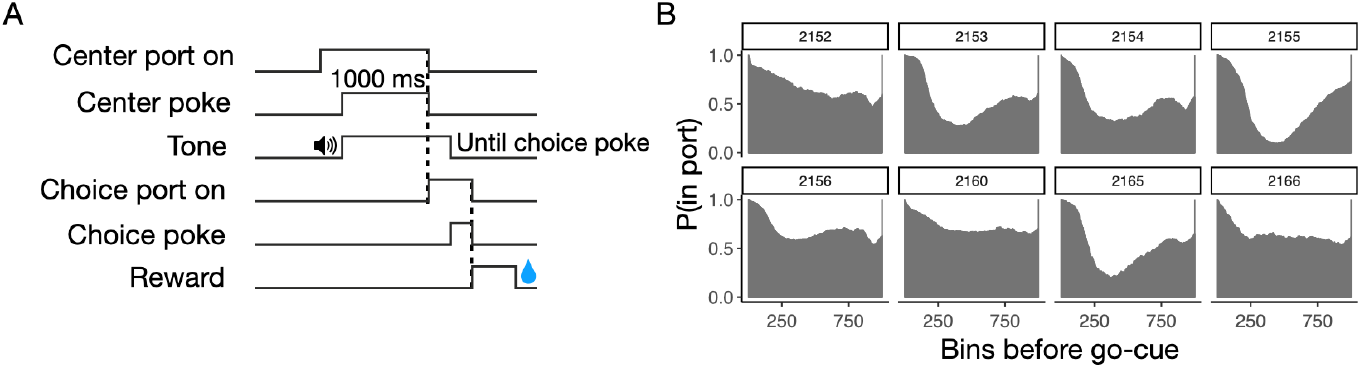
Task and behavior. **A.** Illustration of the timeline of a risky choice trial. **B.** The probability of being in the center port before the go-cue and after the first center port poke. For each trial, the period between the initial poke and the go-cue was segmented into 1000 bins. Go-cue is defined as the onset of the choice port lights. For each bin, a binary value was obtained to indicate whether the animal was in the port or not. The probability of being in the port for each bin was calculated by taking the mean of the binary vector. Only control trials were used for this analysis.

**Figure S2.**
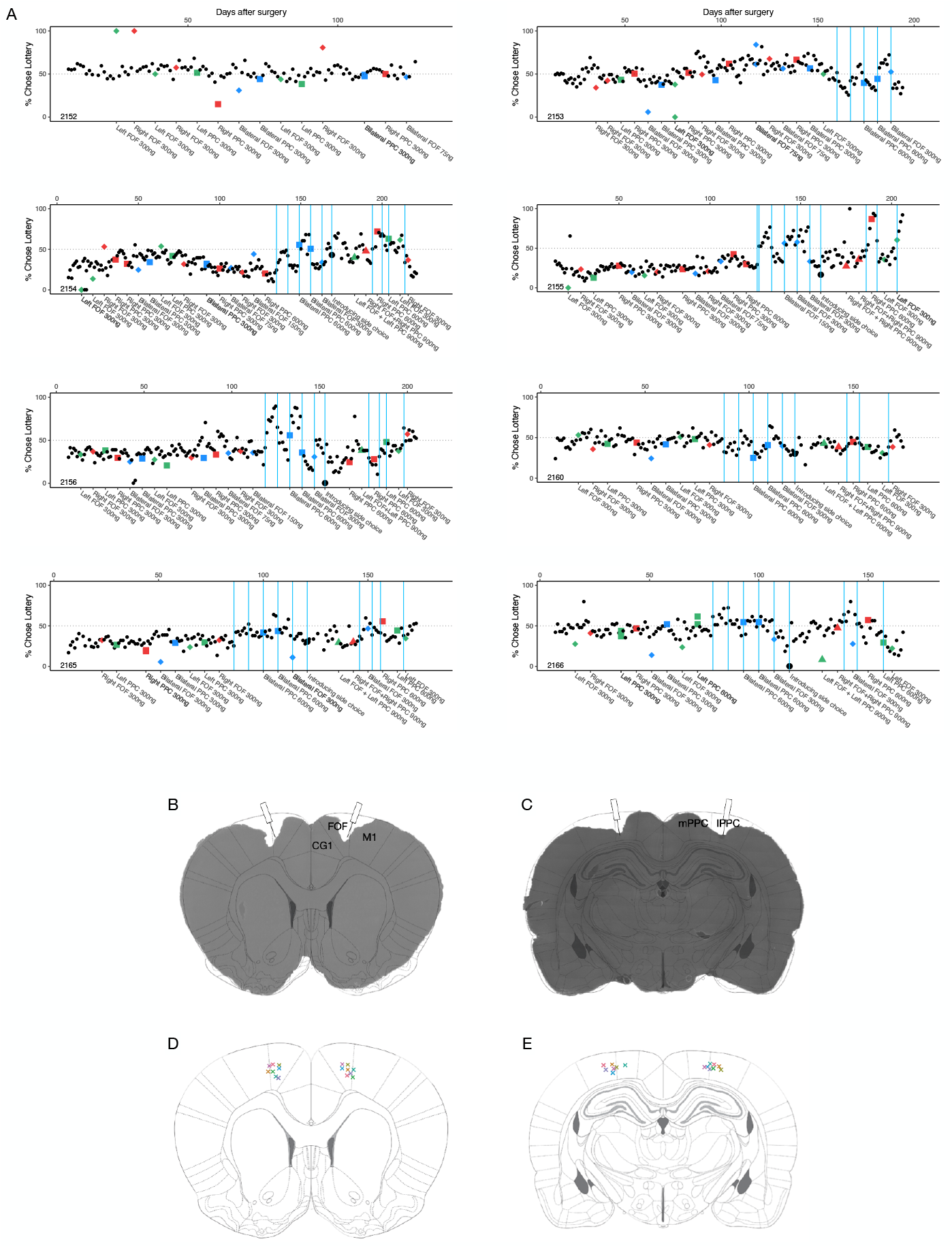
**A**. Timeline of percentage choosing the lottery in each session for each rat and a visual summary of the experimental treatments. Each point is the percentage choosing lottery for the given session. The number at the x-axis indicates the days passed since the surgical implantation of cannulae. Control days are shown as small black dots. Right infusions are shown in red, left infusions are in green, and bilateral infusions are shown in blue. FOF infusions are represented by diamonds, PPC infusions by squares, both FOF and PPC infusions by triangles. The blue bars indicate the day of a model-based surebet value change. The large black dot indicates the day when free choice trials were introduced. The bottom x-labels describe the details (side, region and dose) of each infusion. **B.** Coronal section of an example rat brain showing cannulae implanted at 20° in FOF, overlaid with a section 2.04 mm anterior to Bregma (Paxinos and Watson, 2004). Note, that in the nomenclature of Paxinos and Watson (2004) the area that we describe as the FOF is considered to be part of M2. CG1 = Cingulate Cortex. **C.** Coronal section of an example rat brain showing cannulae implanted at 10° in PPC, overlaid with a section 3.48 mm posterior to Bregma. mPPC = medial PPC, lPPC = lateral PPC. **D.** Actual cannulae placements in FOF, color represents the subject ID as in Figure 1E. **E.** Cannulae placements in PPC.

**Figure S3.**
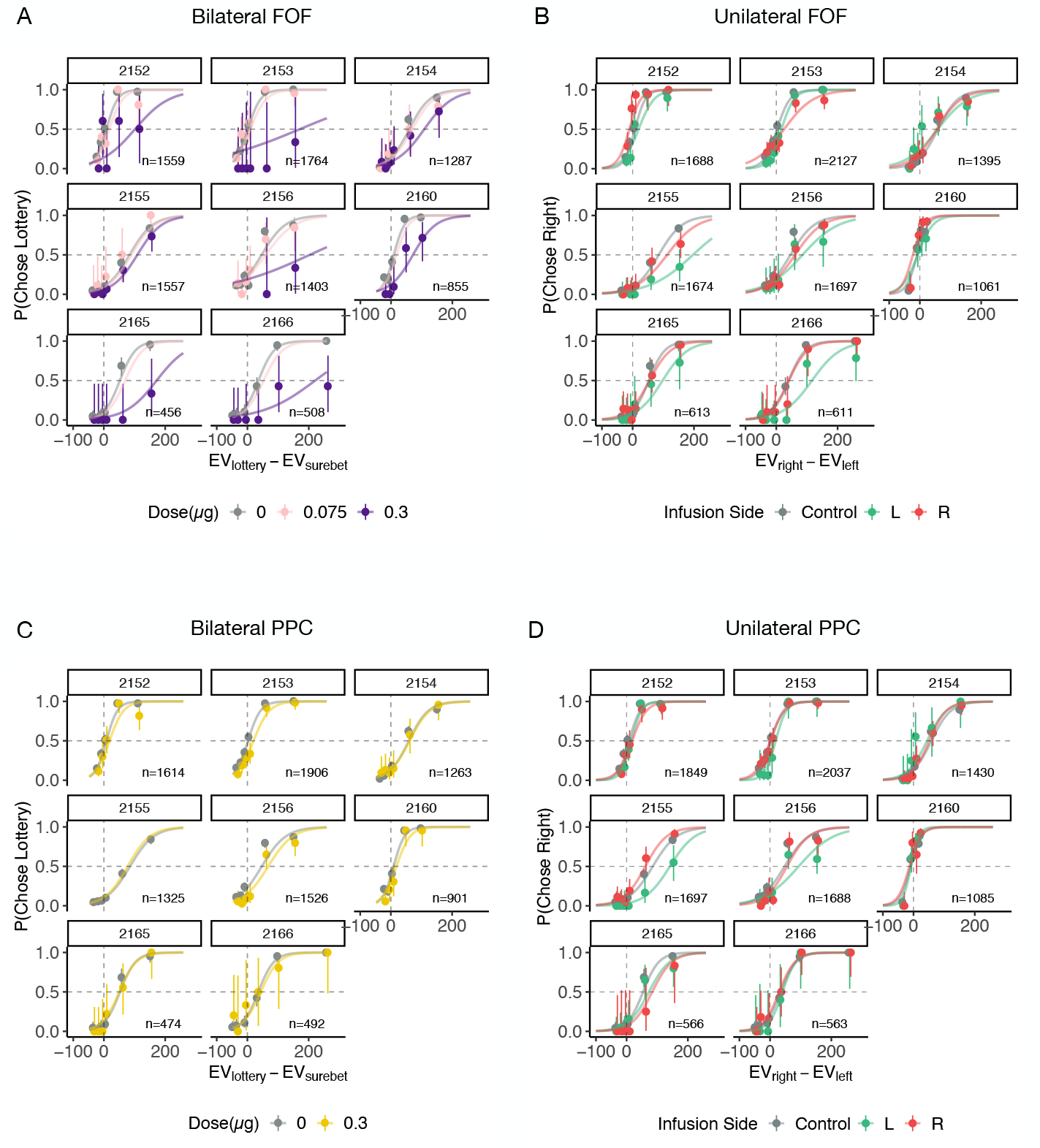
Inactivations by subject. The circles with error bars are the binned mean and 95% binomial confidence intervals. The lines are the model predictions generated by the GLMM. **A.** Bilateral FOF inactivation with 0.075 *μ*g and 0.3 *μ*g muscimol per side. **B.** Unilateral FOF inactivation with 0.3 *μ*g muscimol. **C.** Bilateral PPC inactivation with 0.3 *μ*g muscimol per side. **D.** Unilateral PPC inactivation with 0.3 *μ*g muscimol.

**Figure S4.**
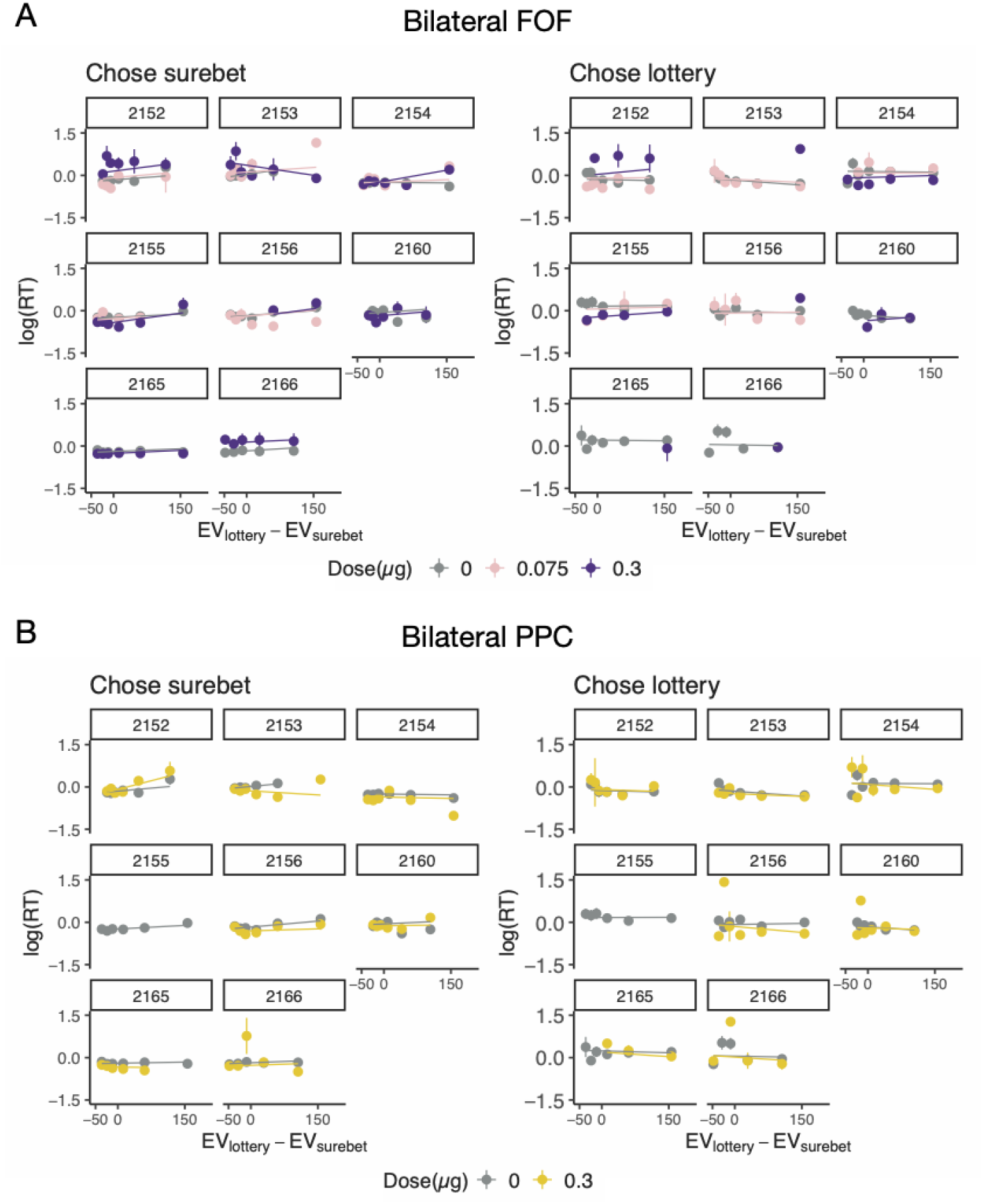
Reaction time (RT) and LMM model fits for bilateral FOF and PPC inactivations. The circles with error bars represent the mean and standard error of log(RT). The lines are the model predictions generated by the LMM. **A.** Bilateral FOF inactivation. Reaction times are from the same trials as presented in Figure S3A. **B.** Bilateral PPC inactivation. Reaction times are from the same trials as presented in Figure S3C.

**Figure S5.**
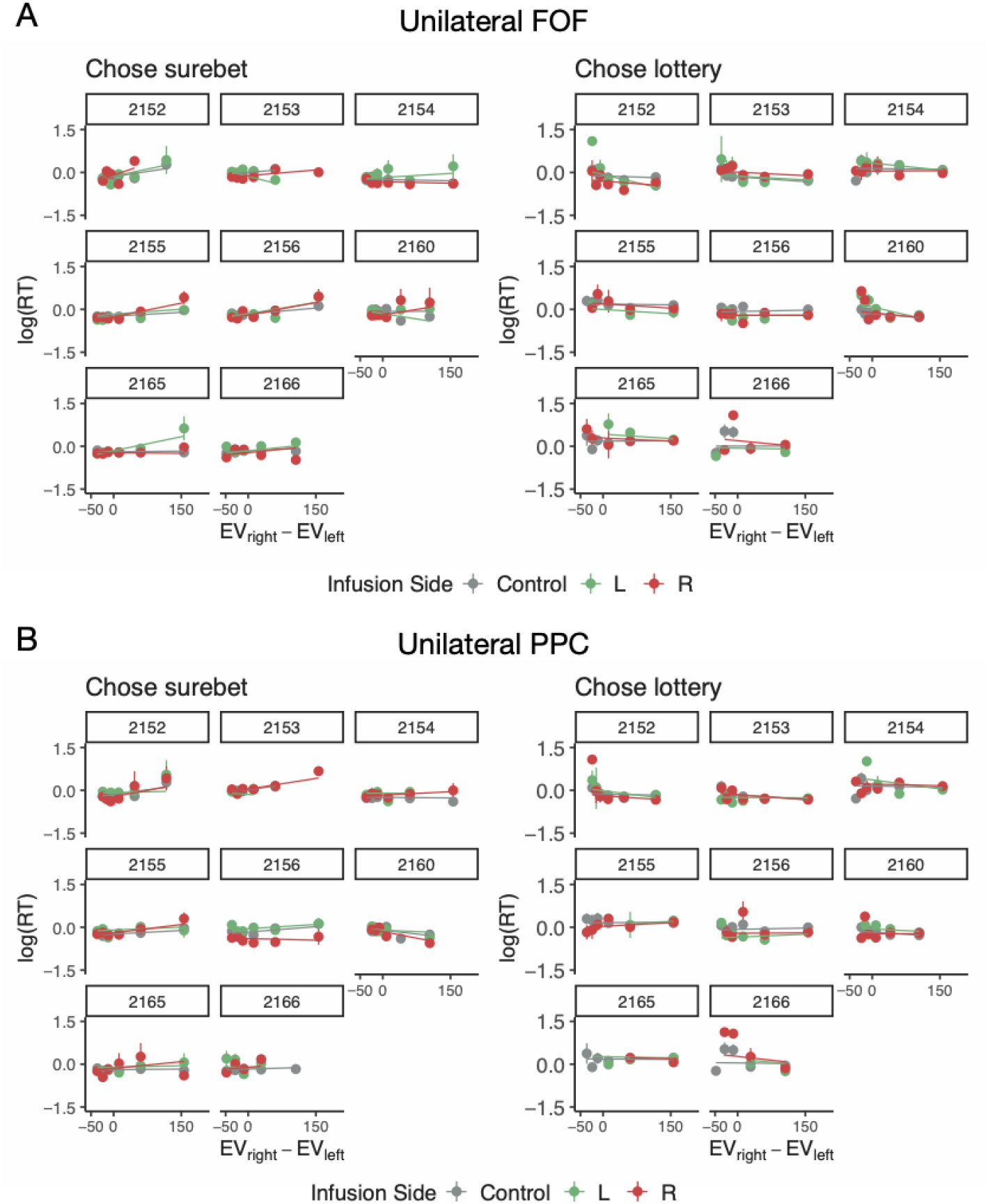
RT and LMM model fits for unilateral FOF and PPC inactivation trials. The circles with error bars represent the mean and standard error of log(RT). The lines are the model predictions generated by the LMM. **A.** Bilateral FOF inactivation. Reaction times are from the same trials as presented in Figure S3B. **B.** Bilateral PPC inactivation. Reaction times are from the same trials as presented in Figure S3D.

**Figure S6.**
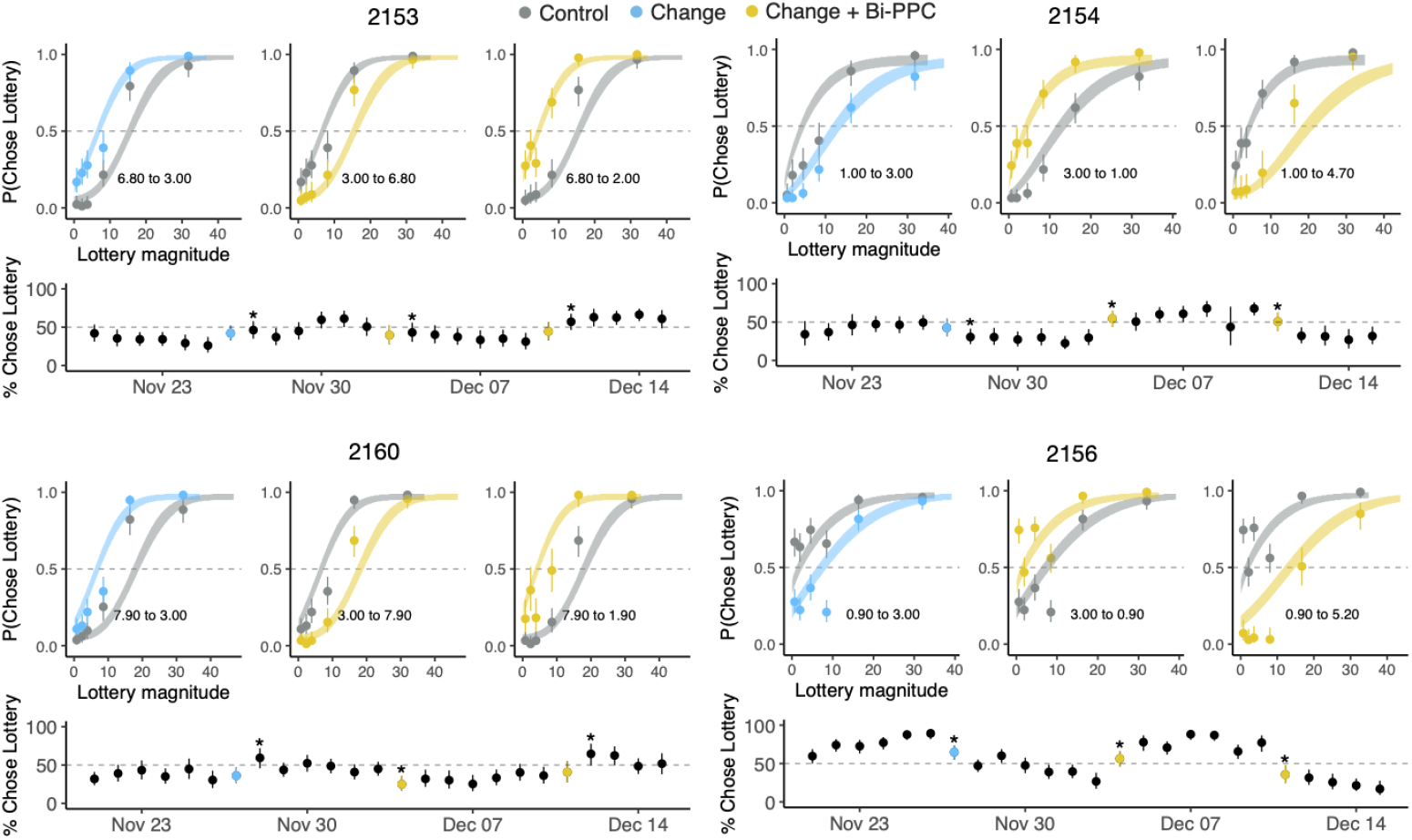
Behavioral adaptation of subject 2153, 2154, 2156 and 2160 in the surebet learning experiment. Only one model was fit to all the trials and used for prediction for each animal. *Top three subpanels*: the circles with error bars are the binned mean and 95% binomial confidence intervals; the ribbons are generated using the fit parameter posterior of with 80% confidence intervals. Behavior from 6 sessions immediately before a surebet change is in gray, behavior from 7 sessions after a surebet change (including the very day) is in light blue if no infusion, in gold if with 0.6 *μ*g bilateral PPC infusion. Text annotation shows the old and new surebet magnitudes. *Bottom subpanel*: The percentage choosing lottery of each session. Asterisk indicates when change in choices can be significantly detected on that session compared to the previous 6 sessions with old surebet magnitude.

**Figure S7.**
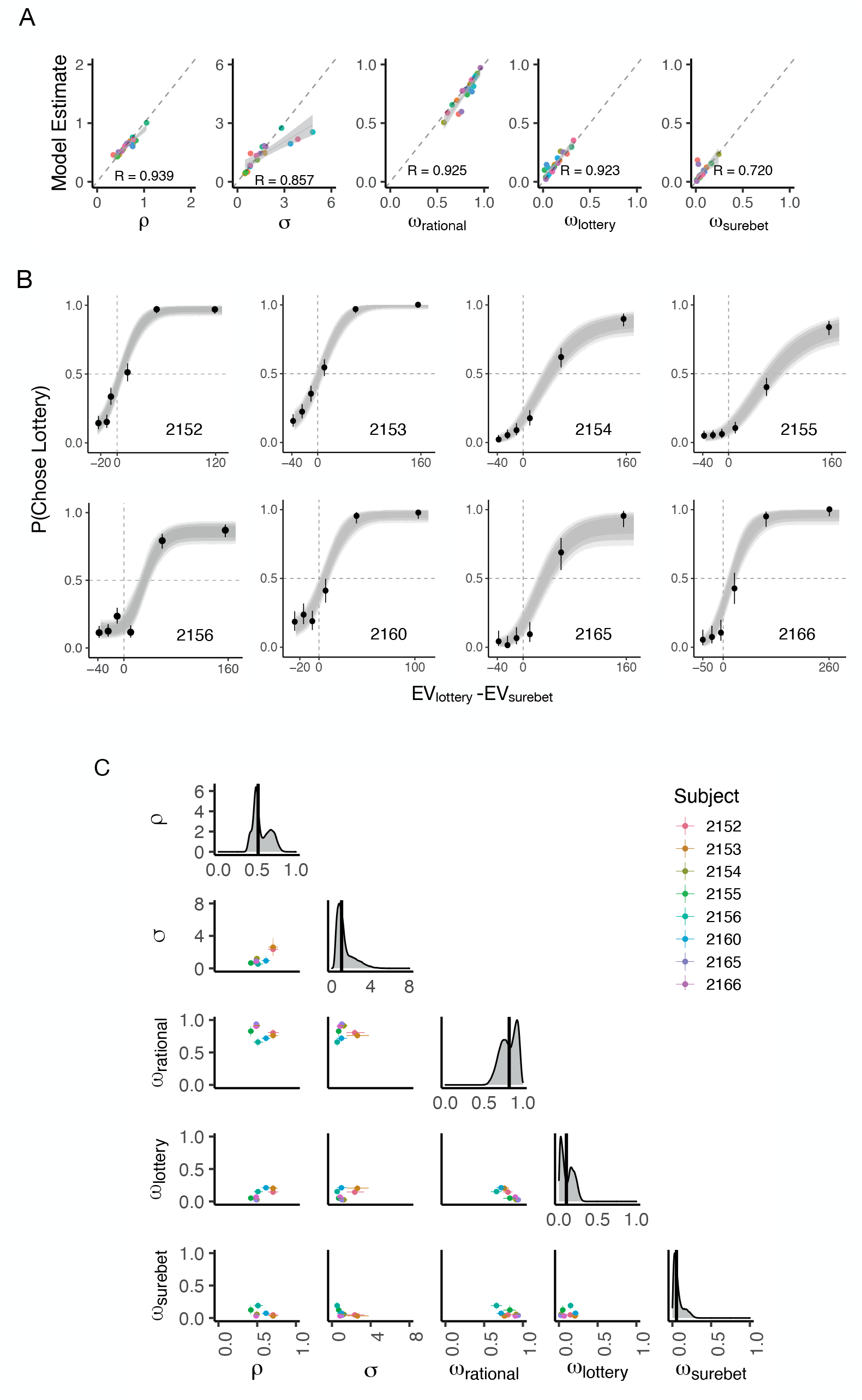
The three-agent mixture model. **A.** The model can recover the data-generating parameters well. Twenty Synthetic datasets were created by sampling from the same prior distributions as specified in Methods. The true parameter value is on the x-axis, the maximum a posteriori model estimation is on the y-axis. Color represents the identity of each synthetic dataset. All the parameters fall along the diagonal line. **B.** The psychometric data and model prediction from the three-agent mixture model for 8 animals. The circles with error bars are the binned mean and 95% binomial confidence intervals. The ribbons are model predictions generated using the fitted parameters. The dark, medium and light shade represent 80%, 95% and 99% confidence intervals, respectively. Data used are the same as the control sessions in Figure 1C. **C.** Summary of the fit model parameters from the control sessions of 8 animals. The mean and 95% confidence interval of each parameter pair are shown in the off-diagonal, colored by subject. Density plots of concatenated posterior samples for each parameter are on the diagonal, the black bar denotes the median.

**Figure S8.**
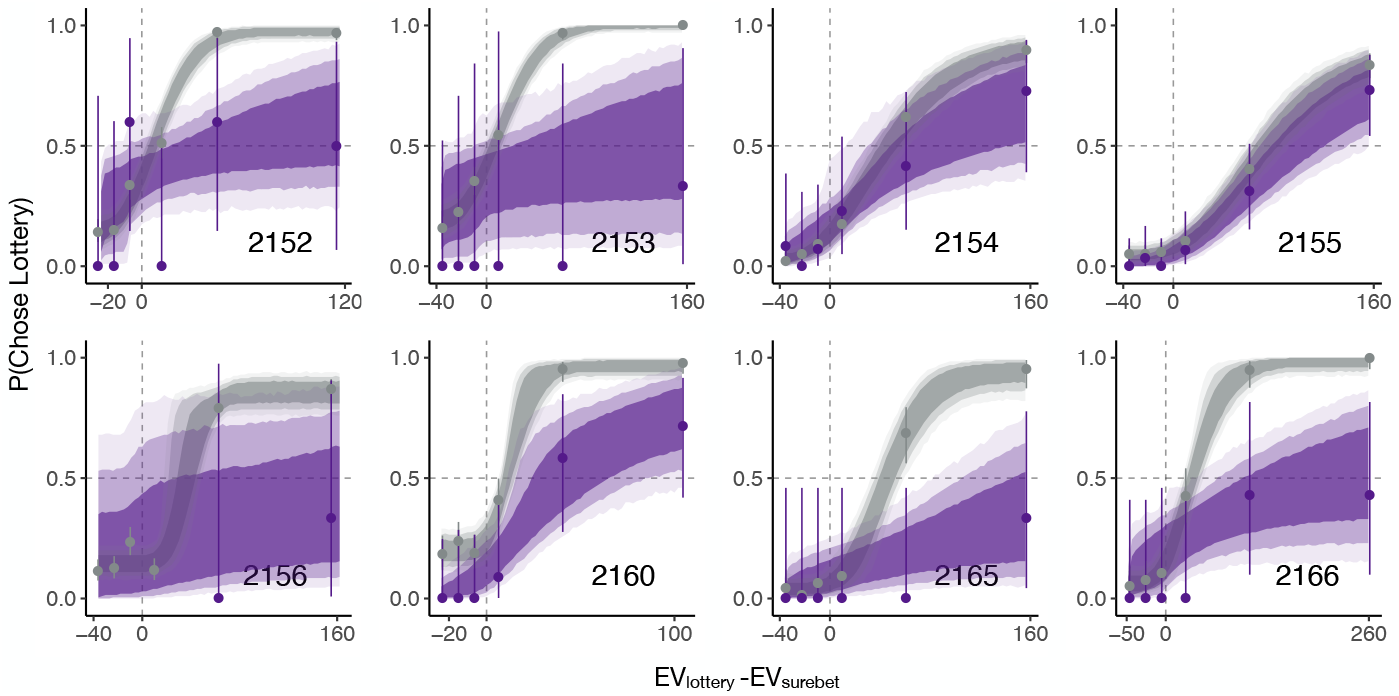
Subjects’ choices superimposed with the inactivation model fit on control (in gray) and bilateral FOF inactivation (in purple) dataset simultaneously. The circles with error bars are the binned mean and 95% binomial confidence intervals. The ribbons are model predictions generated using the fitted parameters. The dark, medium and light shade represent 80%, 95% and 99% confidence intervals, respectively.

**Figure S9.**
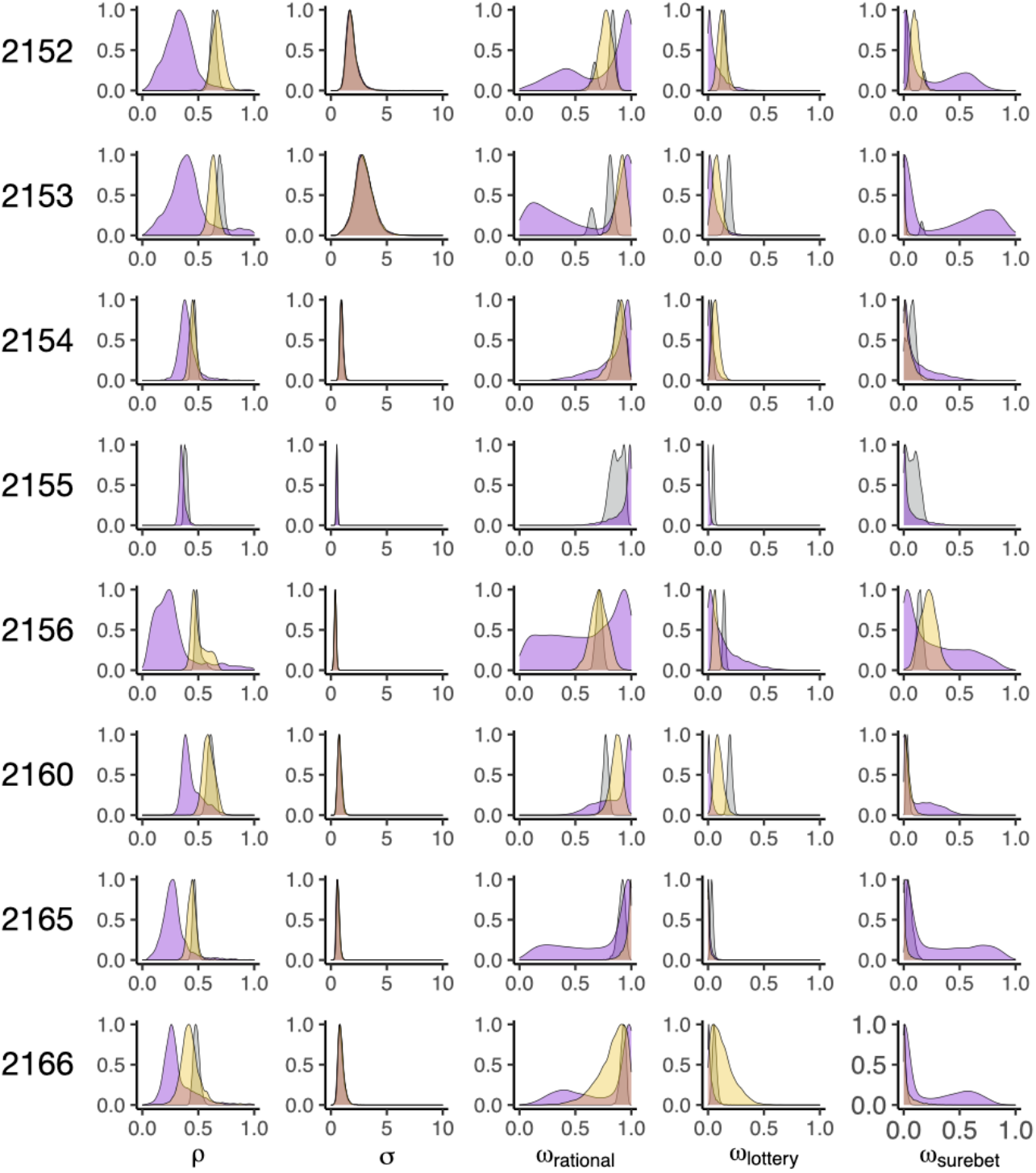
Posterior distributions for each parameter using the inactivation model fit to the control (in gray), 0.3 *μ*g per side bilateral FOF inactivation (in purple) and 0.3 *μ*g per side bilateral PPC inactivation (in gold) dataset simultaneously. To allow easier visual comparison, all posteriors were normalized so that the peak of the distribution was set to 1. Since subject 2155 lost one PPC cannula, only the control and bilateral FOF fit was included here. From left to right: *ρ* is the exponent on the utility function, *σ* denotes the noise in utility representation, *ω_rational_* is the weight of the rational agent, *ω*_lottery_ is the weight of the lottery agent, and *ω_surebet_* is the weight of the surebet agent.

**Figure S10.**
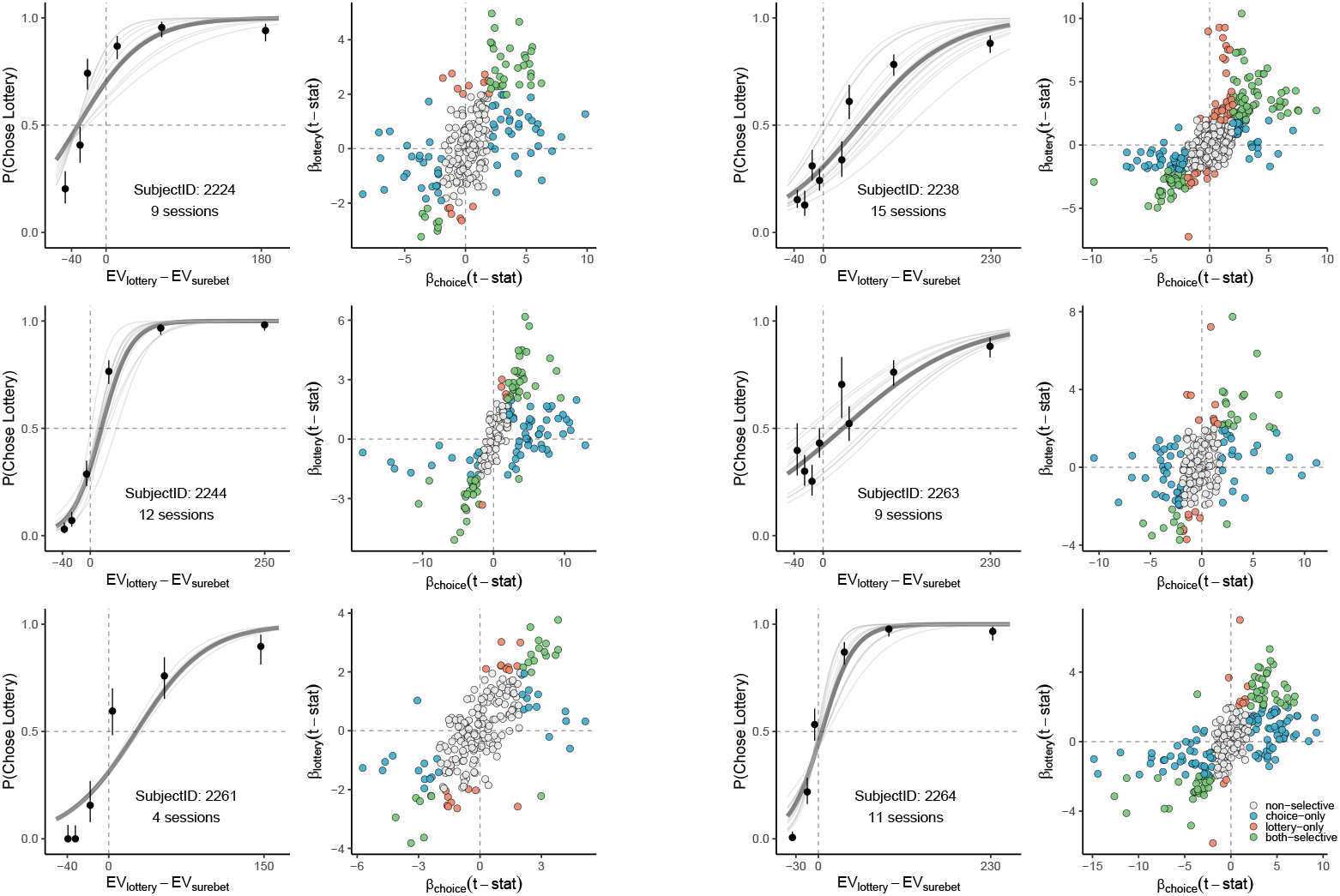
Behavior and the distribution of coefficients for the up-coming choice and lottery value based on the mixed effects linear models for each individual animal. For each subject, the left panel shows the individual’s behavioral performance for all the electrophysiology recording sessions. The dots with error bars show the probability of choosing lotteries plotted against Δ expected value of the two options. The lines are the psychometric curves estimated by a logistic fit to the data, the thin gray lines are fit to each session, the thick gray line fit to all the sessions combined. The right panel shows the distribution of coefficients for the lottery value and upcoming choice for all the neurons recorded in each animal. Gray dots indicate the non-task relevant neurons, light blue dots indicate the purely choice neurons, orange dots indicate the purely lottery selectivity neurons, and green dots indicate tuning for both upcoming choice and lottery values.

